# SATB1 is a targetable modulator of JAK-STAT signaling and cytokines in human Treg and Tconv cells

**DOI:** 10.64898/2026.02.13.705474

**Authors:** Saskia Kolb, Leonie Diekmann, Elizabeth D. Lochert, Linda Warmuth, Julia Ritter, Michael Weber, Markus Hoffmann, Markus List, Daniel Kotlarz, Isabelle Serr, Carolin Daniel, Dirk H. Busch, Christian Schmidl, Kathrin Schumann

## Abstract

The chromatin organizer SATB1 is indispensable for thymic regulatory T cell (Treg cell) development and T helper cell induction. Several gene loci have been described to be SATB1-controlled, including the transcription factor GATA3 and the cytokine loci IL-4 and IL-17. However, the global effects of SATB1 on fully differentiated human CD4 conventional T cells (Tconv cells) and Treg cells, and thus SATB1’s potential as a target for T cell engineering, are poorly understood. We describe SATB1-regulated gene signatures as largely subset-specific, with broader effects on Treg cells. Despite of the distinct gene-regulatory patterns, we observe overarching dysregulated cytokine and JAK-STAT signaling after *SATB1* ablation. Functionally, *SATB1* KO reduces human Treg cell suppressive capacities but boosts tumor clearance via CD4 CAR T cells in a preclinical, humanized mouse model. Together, Treg destabilization and simultaneous increased activation of CD4 CAR T cells by SATB1 modulation may be an interesting strategy to boost the efficiency of CAR T cell therapies.

## INTRODUCTION

The impact of Satb1 (SATB Homeobox 1), a transcription factor and chromatin organizer, on T cell function, has been first described in KO mouse models. *Satb1* knockout (KO) mice have a reduced overall survival as well as small thymi and spleens as a result of arrested T cell development (Alvarez *et al*, 2000). During thymic development, Satb1 is induced in a TCR-dependent manner and, in turn, regulates the expression of lineage-defining genes such as *Runx3*, *Cd8*, *Cd4* as well as the Treg master transcription factor *Foxp3* (Kakugawa *et al*, 2017; Kitagawa *et al*, 2017). Mice with conditional *SATB1* KO in CD4 T cells have lower Foxp3+ Treg cell numbers and reduced DNA hypomethylation at *Foxp3* conserved non-coding region 2 (CNS2), indicating cellular destabilization (Kitagawa *et al*., 2017).

However, our knowledge of Satb1/SATB1 in fully differentiated human CD4 T cells is still incomplete, particularly during inflammation. Interestingly, in fully differentiated human Treg cells, SATB1 is a positive regulator of FOXP3 expression (Schumann *et al*, 2020). Foxp3/FOXP3 is counteracting this upregulation by inducing microRNAs, which in turn downregulate SATB1 mRNA levels in human and murine Treg cells indicating tight regulation for proper cellular function (Beyer *et al*, 2011). Overexpression of SATB1 in Treg cells results in increased levels of multiple pro-inflammatory cytokines associated with Th1, Th2, and Th17 effector functions, which overall resulted in reduced suppressive capacity (Beyer *et al*., 2011; Burute *et al*, 2012). Also, Tconv cells depend on Satb1 in the steady state for proper cell function. *Satb1* KO Tconv cells have been described to be more susceptible to Treg cell-mediated suppression due to lower expression levels of CD25 and IL-2 (Gupta *et al*, 2022). Satb1 is further known as a regulator of different cytokine loci in Th2 cell differentiation in the periphery and regulator of Th17 differentiation, effector tissue phenotype in experimental autoimmune encephalomyelitis (Ahlfors *et al*, 2010; Köhne *et al*, 2025; Yasuda *et al*, 2019). Transcription factors are increasingly the focus for novel chimeric antigen receptor (CAR) T cell engineering approaches to stabilize or induce certain effector phenotypes (Dai *et al*, 2024; Doan *et al*, 2024). However, SATB1-controlled gene signatures have not been dissected within this context and their functional consequences are unclear.

In this study, we analyzed the impact of SATB1 ablation in fully differentiated CD4 T cells on chromatin and mRNA level and the resulting changes for key signaling pathways and effector functions in a cell-type-specific manner. Analyzing SATB1’s gene and chromatin modulation during pro-inflammatory stimulation reveals largely distinct chromatin and gene signatures in CD4 Tconv and Treg cells, with a greater impact on gene signatures in Treg cells. JAK-STAT and cytokine/cytokine receptor signaling pathways are dysregulated in both cell types in a pro-inflammatory microenvironment after SATB1 ablation. Functional validation of *SATB1* KO cells revealed diminished suppressive capacity in Treg cells and improved effector function in CAR CD4 T cells.

## RESULTS

### SATB1 is a regulator of cytokine and FOXP3 expression in Treg and Tconv cells

To dissect the impact of SATB1 gene regulation in human CD4 T cells in the steady state and during inflammation, we ablated either the safe-harbor locus *AAVS1* as a negative control or the transcription factor *SATB1* using Cas9 ribonucleoprotein nucleofection of *in vitro* expanded Treg and CD4 Tconv cells (Fig. 1A) (Schumann *et al*., 2020). To mimic an inflammatory microenvironment, as observed in inflammatory bowel disease or also in solid tumors, both KO T cell subsets were challenged with high doses of pro-inflammatory IL-12 (Mirlekar & Pylayeva-Gupta, 2021; Verstockt *et al*, 2023). Currently, several strategies to locally administer IL-12 to tumors to boost the efficiency of adoptive T cell therapies are being tested in the clinic (Minnar *et al*, 2023). Interestingly, exposure of Treg cells to high doses of IL-12 can additionally induce a Th1-like, destabilized Treg cell phenotype (Dominguez-Villar *et al*, 2011).

**Figure 1.**
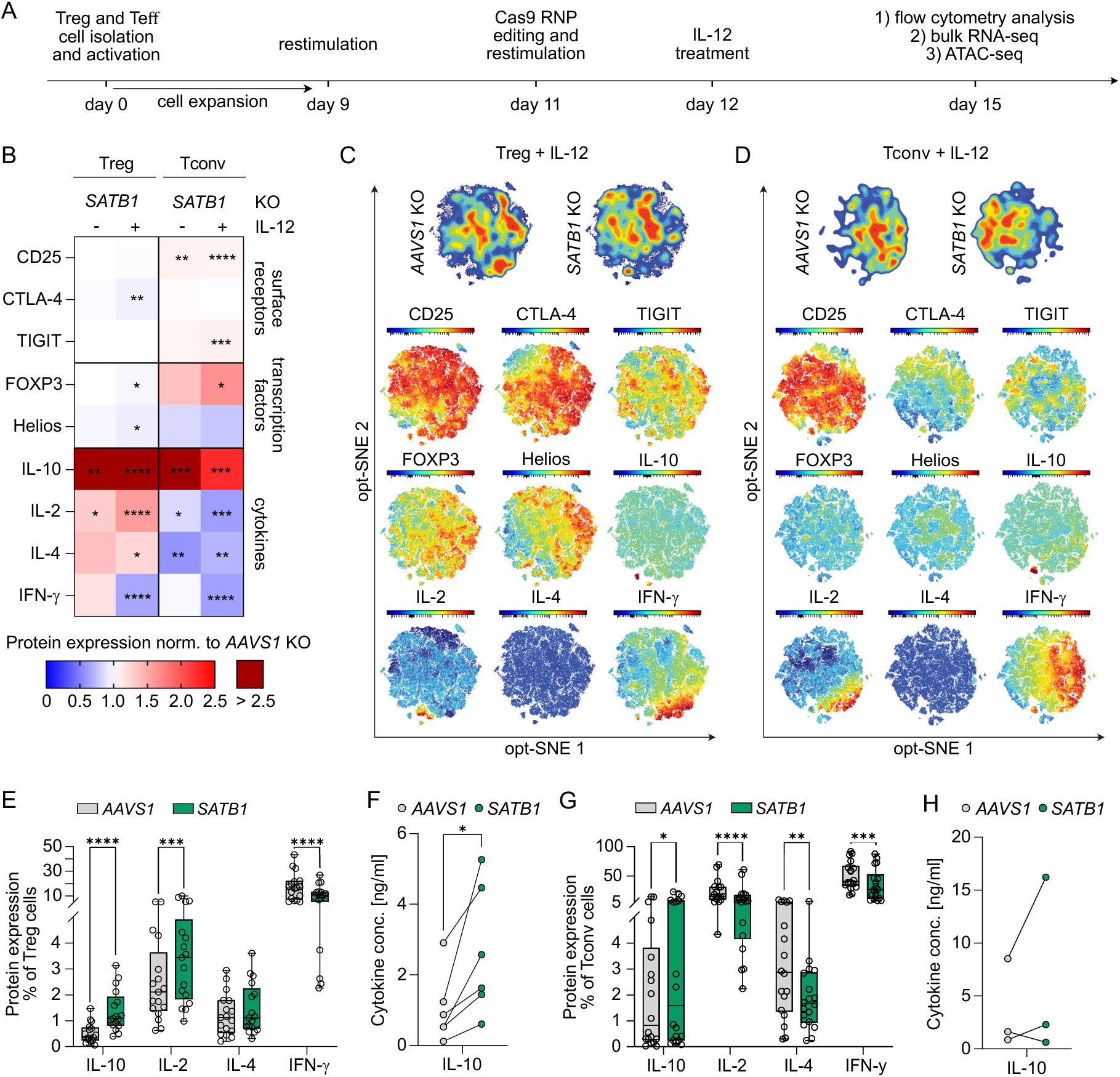
*SATB1* KO causes a destabilized Treg phenotype and enhanced activation in Tconv cells. **(A)** Workflow of *in vitro SATB1* KO experiments in human Treg and Tconv cells. Treg and Tconv cells were isolated from blood of healthy donors, *ex vivo* expanded, CRISPR edited, and challenged with or without IL-12. KO cells were phenotypically characterized by flow cytometry, bulk RNA-seq, and ATAC-seq analysis. **(B)** Heatmap displaying fold changes of pro- and anti-inflammatory flow cytometry markers in *SATB1* KO Treg cells and *SATB1* KO Tconv cells normalized to *AAVS1* KO control cells. The fold change was calculated based on the percentage of the individual marker pre-gated on living cells. n = 12-18, ratio paired t test. **(C, D)** opt-sne contour plot of IL-12-treated *AAVS1* KO and *SATB1* KO Treg **(C)** and Tconv cells **(D)**. Expression levels (MFI) of tested flow cytometry markers plotted on opt-SNE plot, (n = 24). **(E)** Absolute percentages of intracellular cytokine flow cytometry stainings of IL-12 treated *AAVS1* KO and *SATB1* KO Treg cells. n = 12-17, paired t test. **(F)** Extracellular IL-10 levels determined by LEGENDplex^TM^ assay of IL-12-treated *AAVS1* KO and *SATB1* KO Treg cells. n = 6, paired t test. **(G)** Absolute percentages of intracellular cytokine flow cytometry stainings of with IL-12 treated *AAVS1* KO and *SATB1* KO Tconv cells. n = 12-18, paired t test. **(H)** Extracellular IL-10 levels determined by LEGENDplex^TM^ assay of IL-12 treated *AAVS1* KO and *SATB1* KO Tconv. n = 3, paired t test. * p<0.05, ** p<0.01, *** p<0.001, **** p<0.0001

The *AAVS1* and *SATB1* KO rates were comparable in Treg and Tconv cells. IL-12 supplementation reduced the variability in the KO frequencies between donors (Fig. S1A). Successful *SATB1* KO could also be confirmed on the mRNA level with higher basal SATB1 expression in Treg cells as previously described (Fig. S1B) (Beyer *et al*., 2011).

We also included the Treg master transcription factor FOXP3 and the transcriptional regulator Helios, that has been associated with Tconv cell activation and Treg cell stability (Allan *et al*, 2007; Lam *et al*, 2022). Both transcription factors were very slightly decreased after *SATB1* ablation in Treg cells after IL-12 stimulation (Fig. 1B; Fig. S1E; gating strategy: Fig. S1D). SATB1 has been previously described as a positive regulator of FOXP3 expression (Chaurio *et al*, 2022; Kitagawa *et al*., 2017). In our experimental setting, *SATB1* ablation in fully differentiated Treg cells resulted in slight reductions of FOXP3 protein as well as mRNA levels (Fig. 1B; Fig. S1B, E). To still exclude a dominant effect of FOXP3 reduction in *SATB1* KO Treg cells, we included *FOXP3* KO Treg cells as an additional control condition to separate FOXP3-from SATB1-dependent effects. These two KO conditions differed in their cytokine profile as well as surface marker expression (Fig. 1B, Fig. S1C, Fig. S1E). Together these results indicate that in our experimental setting phenotypic changes in *SATB1* KO Treg cells are largely FOXP3-independent. In *SATB1* KO Tconv, the expression of FOXP3 was heightened whereas Helios was reduced (Fig. 1B; Fig. S1B, G).

Phenotypic changes in KO cell stability were assessed by flow cytometry staining of pro- and anti-inflammatory protein markers (Fig. 1B; Fig. S1E, F; gating strategy: Fig. S1D). Slight, yet reproducible, changes were observed for the cell activation-dependent surface markers CD25, CTLA-4, and TIGIT. For example, *SATB1* KO Tconv cells upregulated CD25 and TIGIT especially after IL-12 treatment (Fig. 1B, Fig. S1G). To analyze the impact of these more subtle changes we plotted the data of IL-12 conditioned cells on opt-SNE density plots. *SATB1* KO Treg and Tconv cells showed clear shifts compared to their *AAVS1* KO counterparts (Fig. 1C, D). Areas enriched for *SATB1* KO Treg cells were defined by lower levels of FOXP3, CTLA-4 and Helios, and higher IL-2 (Fig. 1C). *SATB1* KO Tconv cells depicted higher levels of TIGIT and reduced IL-2 and IFNγ (Fig. 1D). Similar patterns could be detected in control-treated cells. However, the differences in control untreated *SATB1* KO Treg cells were less pronounced compared to IL-12 conditioning (Fig. S2).

Next, we analyzed the cytokine profile of *SATB1* KO Treg and Tconv cells in more depth. *SATB1* KO Treg cells increased their intracellular production of the pro-inflammatory cytokines IL-2 and IL-4 with or without IL-12 conditioning and IFNγ without IL-12 treatment (Fig. 1B, C, E, Fig. S1E). *SATB1* ablation in Tconv reduced overall their proinflammatory cytokine production (Fig. 1B, D, G, Fig. S1G). An increase in IL-10 secretion was observed in *SATB*1 KO Treg as well as Tconv cells based on flow cytometry (Fig. 1B-E, G; Fig. S1E, G). High IL-10 levels have been correlated with heightened Tconv cell activation and Treg cell suppressive capacity (Sun *et al*, 2009). These changes in IL-10 production were confirmed for *SATB1* KO Treg cells by extracellular detection with LEGENDplex assay (Fig. 1F; Fig. S1F). In general, taking all the tested markers into account, IL-12 treatment resulted in less phenotypic donor variation and higher statistical significance in most flow cytometry markers tested (Fig. 1B).

Overall, *SATB1* ablation in Treg and Tconv cells resulted in subset-specific regulation of the pro-inflammatory cytokines and a general boost in IL-10 production.

### SATB1-regulated gene expression and chromatin accessibility are largely distinct in Treg and Tconv cells

Our flow cytometry data showed distinct shifts in proinflammatory marker expression in *SATB1* KO Treg and Tconv cells. To test whether these subset-specific gene regulatory patterns extend to further pathways, *AAVS1* and *SATB1* KO cells were subjected to ATAC-seq and RNA-seq analysis. As we observed more statistically significant changes in protein marker expression after IL-12 stimulation, we focused on KO cells exposed to this pro-inflammatory stimulus (Fig. 1B). Further, we previously identified exposure to a pro-inflammatory environment as an efficient strategy for the dissection of gene networks regulated by individual transcription factors (Schumann *et al*., 2020).

ATAC-seq data revealed a larger number of differently regulated peaks in *SATB1* KO Treg cells (28,363) compared to *SATB1* KO Tconv cells (17,142). 10,610 of these chromatin peaks were regulated in both *SATB1* KO T cell subsets (Fig. 2A). These widespread changes in chromatin accessibility separated the two T cell subsets on principal component 1 in the PCA (PCA plot: 73% variance) but also discriminated *SATB1* KO in comparison to *AAVS1* control cells on PC2 (PCA plot: 21% variance) (Fig. 2B). Analysis of RNA-seq data of *SATB1* KO Treg and Tconv cells revealed 4 times more dysregulated genes in Treg cells (2,479 genes) compared to Tconv cells (628 genes) (Fig. 2C), which resulted in strong separation of the subsets (PCA: 92% variance) (Fig. 2D). Only 108 mRNA changes were conserved between both T cell subsets (Fig. 2C, E). The majority of these mRNA changes were co-regulated, meaning either up- or downregulated in *SATB1* KO Treg and Tconv cells. Several transcription factors are part of these co-regulated genes, including the known SATB1-target GATA3 (Fig. 2E). Reduced GATA3 levels in Tconv cells as well as Treg cells after SATB1 ablation with or without IL-12 treatment could also be confirmed on the protein level via flow cytometry (Fig. S3A). In murine *Satb1* KO T cells, a shift from Th2 towards Th1 differentiation has been observed, which aligns with our finding in human Tconv cells (Ahlfors *et al*., 2010; Burute *et al*., 2012). Overall, these data show largely distinct SATB1-regulated gene signatures in Treg and Tconv cells.

**Figure 2.**
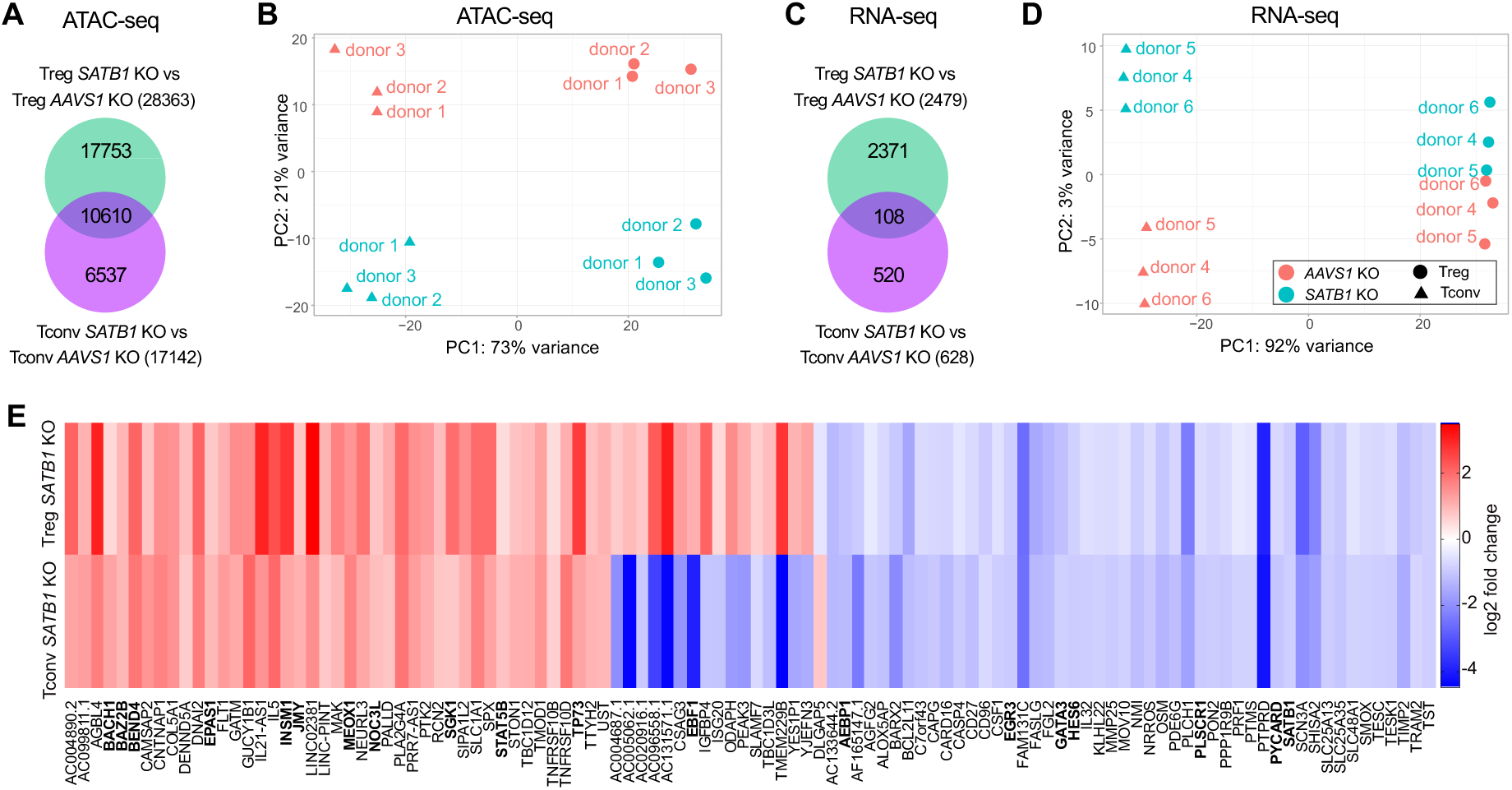
SATB1 controls largely distinct gene signatures in human Treg and Tconv cells after IL-12 treatment. **(A)** Venn diagram of differentially accessible chromatin regions in *SATB1* KO Treg and Tconv cells normalized to AAVS1 KO control cells (p-value < 0.05). a, b: n = 3. **(B)** PCA plot of *AASV1* and *SATB1* KO Treg and Tconv cells treated with IL-12 analyzed by ATAC-seq. **(C)** Venn diagram of differentially expressed genes in *SATB1* KO Treg and Tconv cells normalized to *AAVS1* KO control cells (p-value < 0.05). **(D)** PCA plot of *AASV1* and *SATB1* KO Treg and Tconv cells analyzed by RNA-seq after IL-12 conditioning. **(E)** Heatmaps indicating log2 fold change of overlapping RNA-seq data of *SATB1* KO Treg and Tconv cells treated with IL-12. Human transcription factors were indicated in bold. c – e: n = 3.

### SATB1 is a transcriptional regulator of JAK-STAT signaling in CD4 T cells

Next, we examined transcriptional and chromatin changes in Treg and Tconv KO cells individually. In both T cell subsets, more open chromatin regions after *SATB1* ablation could be detected (Fig. 3A, B). However, the RNA-seq data shows a more complex pattern: In *SATB1* KO Treg cells, similar numbers of genes were up- or downregulated, whereas in *SATB1* KO Tconv cells, the majority of differentially regulated genes were downregulated (Fig. 3C, D).

**Figure 3.**
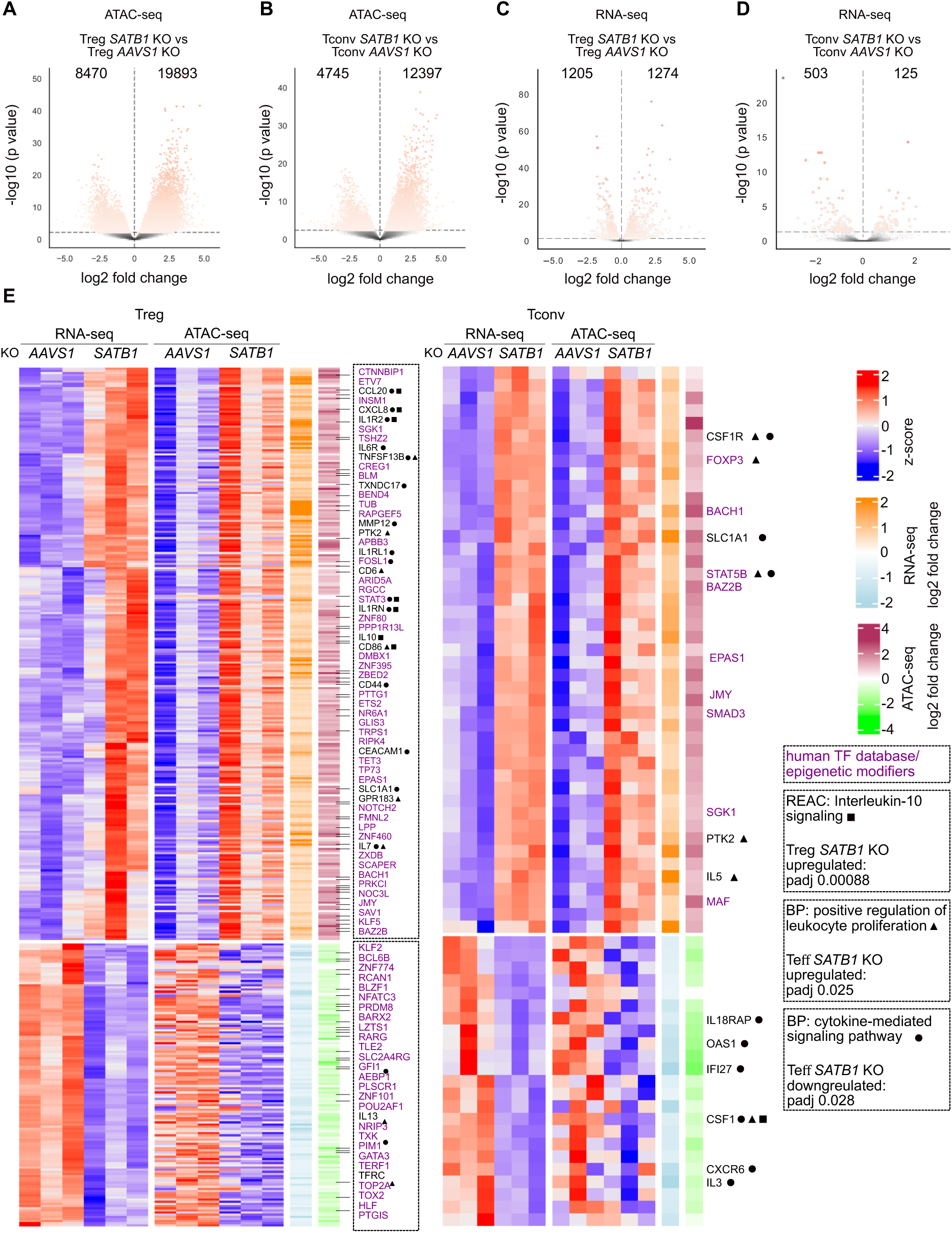
SATB1 differentially controls TF, cytokine expression and proliferation in human *SATB1* KO Treg and Tconv cells after IL-12 treatment. **(A, B)** Volcano plot of differentially accessible chromatin regions of *SATB1* KO Treg versus *AAVS1* KO Treg cells (**A**) and *SATB1* KO Tconv versus *AAVS1* KO Tconv cells (**B**), n = 3. **(C, D)** Volcano plot of differentially expressed genes of *SATB1* KO Treg versus *AAVS1* KO Treg cells (**C**) and *SATB1* KO Tconv versus *AAVS1* KO Tconv cells (**D**). **(E)** Heatmaps displaying z-scores of equally regulated chromatin and gene expression changes in *SATB1* KO Treg and Tconv cells treated with IL-12 normalized to the respective AAVS1 KO controls. Differentially regulated TFs are highlighted in purple. Top hits of Reactome (REAC) and biological pathway (BP) analysis and associated adjusted p-values (padj) if significant are highlighted.

To integrate RNA-seq and ATAC-seq data, changes in chromatin accessibility in a ± 10 kb window around the transcriptional start site were quantified and integrated with changes in mRNA levels to identify subset-specific signatures (Fig. 3E, Fig. S4) (Thakore *et al*, 2024). In Fig. 3E, genes that are regulated in the same manner on chromatin and mRNA level, are depicted, opposite regulation, which was less frequent, in Fig. S4. We identified 267 genes in Treg cells and 45 genes in Tconv cells, which had more open chromatin ± 10 kb around the TSS, and correlated with simultaneous mRNA upregulation after *SATB1* ablation. Accordingly, 132 genes in *SATB1* KO Treg cells and 23 genes in *SATB1* KO Tconv cells had more closed chromatin and were downregulated (Fig. 3E).

Pathway enrichment analysis of the co-regulated gene sets shown in Fig. 3E elucidated a dysregulation of different cytokine secretion and cytokine signaling in Treg and Tconv cells (full list of pathways: Table S6). Interestingly, in *SATB1* KO Tconv cells, the signature “positive regulation of lymphocyte proliferation” (Biological Pathways (BP)) was enhanced, which is in line with the upregulation of FOXP3, TIGIT and IL-10 (Fig. 1B, D, G, H). The mentioned BP pathway also affected the oppositely regulated genes in the ATAC-seq and RNA-seq data sets. However, these changes were not significant (Fig. S4).

In *SATB1* KO Treg cells, the IL-10 signaling-related genes, such as *IL10*, *CD86*, and *IL7*, were significantly upregulated (REACTOME database, term “Interleukin 10 signaling”, Table S6). Upregulation of CD86 expression after deletion of SATB1 in Treg cells could be confirmed on the protein level (Fig. S3B). In *SATB1* KO Tconv cells, genes involved in cytokine-mediated signaling, such as *IL5*, were upregulated, whereas others, including *IL3*, were downregulated (Fig. 3E; BP “cytokine-mediated signaling pathway”, Table S6). Next, we performed KEGG analysis on the global RNA-seq results of *SATB1* KO Treg and Tconv cells to further dissect these patterns. We observed a strong dysregulation of genes related to KEGG “cytokine-cytokine receptor interactions” with more pronounced effects in Treg cells compared to Tconv cells. Again, changes in *SATB1* KO Treg cells were more prominent. Cytokines and cytokine receptors were mostly upregulated (e.g. IL-6, IL-10, IL-5, IL7R, IL1LR2, IL6R). Chemokines and chemokine receptors showed a more diverse up- and downregulation. In Tconv cells overall less genes of this pathway were differentially regulated and their expression was mostly reduced (Fig. 4A).

**Figure 4.**
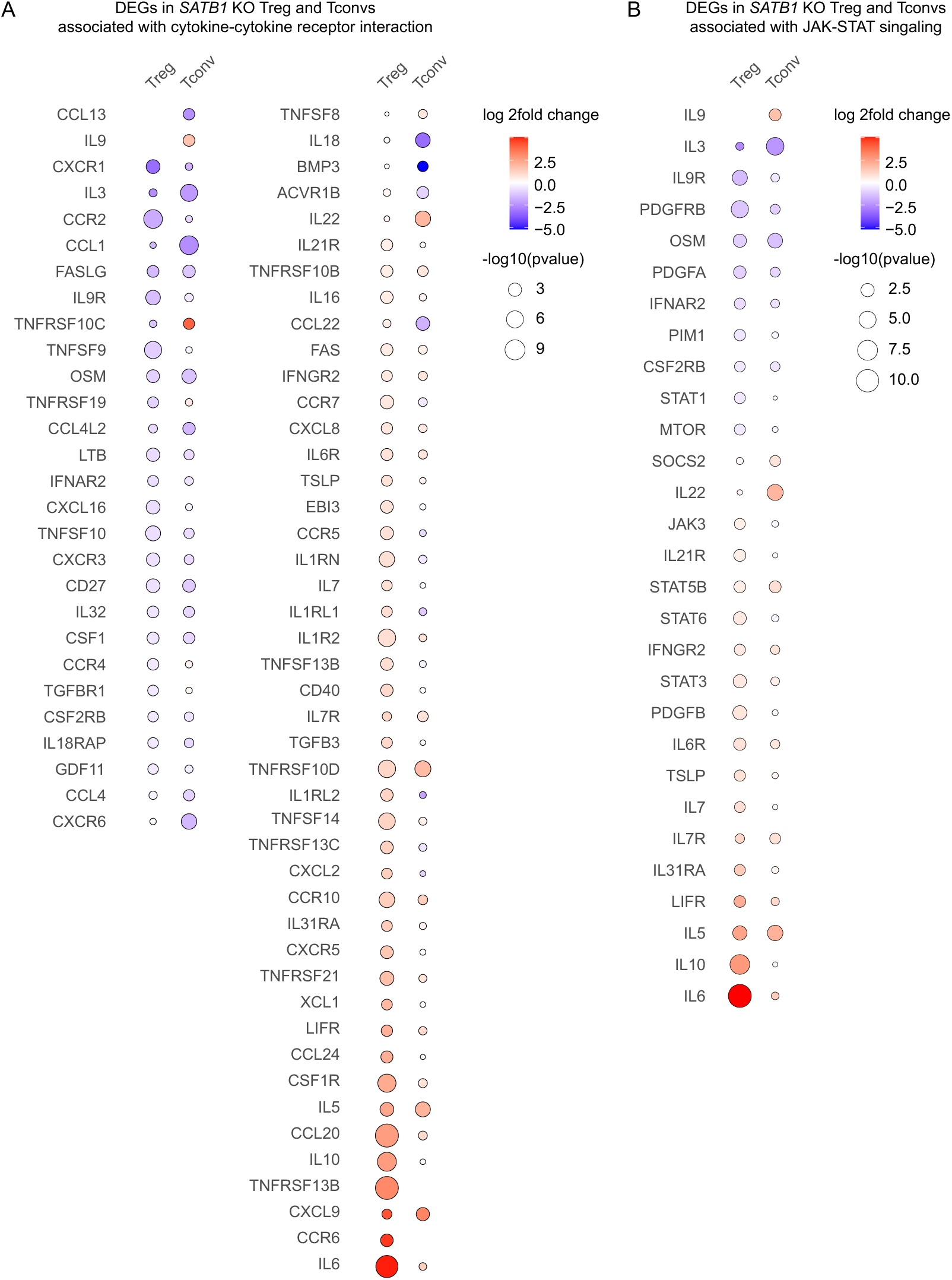
SATB1 differentially regulates cytokine expression and JAK-STAT signaling in human Treg and Tconv cells. Differently expressed genes (DEGs) of the cytokine-cytokine receptor pathway **(A)** and the JAK-STAT signaling pathway **(B)** in *SATB1* KO Treg and Tconv cells treated with IL-12 normalized to the respective IL-12 *AAVS1* KO conditions based on RNA-seq data. Color scheme indicated log2 fold changes of gene expression and the size of the dots the -log10 of the p-value.

Besides that, the *SATB1* KO resulted in a massive dysregulation of transcription factors and epigenetic modifiers in Treg cells and, to a lower extent, in Tconv cells (Fig. 2E, Fig. 3E, Fig. S4). STAT3, a transcription factor with known functions in Treg cell stability as well as IL-10 signaling, was upregulated in *SATB1* KO Treg cells (Aqel *et al*, 2021; Laurence *et al*, 2012; Poholek *et al*, 2016). In *SATB1* KO Tconv cells, the STAT family member STAT5b was significantly more accessible in ATAC-seq, which correlated with higher STAT5b mRNA levels (Fig. 3E). KEGG pathway analysis showed significant changes in “JAK-STAT-signaling”, again more pronounced in *SATB1* KO Treg cells compared to Tconv cells. In *SATB1* KO Treg cells, transcript levels of JAK3, STAT3, STAT5B, and STAT6 were upregulated, and STAT1 was reduced, among other changes. Also, in *SATB1* KO Tconv cellschanges in the JAK-STAT pathway could be observed for example by increased transcript levels of STAT5B and SOCS2 (Fig. 4B).

These results identify SATB1 as a major regulator of the JAK-STAT pathway, cytokine expression, and signaling in fully differentiated human CD4 T cells in a subset-specific manner in a pro-inflammatory microenvironment.

### *SATB1* KO impairs Treg cell suppressive function

So far, flow cytometry, RNA-seq, and ATAC-seq data analysis depicted a complex pattern of gene dysregulation in *SATB1* KO Treg cells affecting a wide range of cytokine/JAK-STAT proteins with an unclear overall impact on Treg cell functionality. For example, STAT3 has been described as a positive regulator of FOXP3 expression in Treg cells, but also as a driver of Treg cell instability (Laurence *et al*., 2012; Pallandre *et al*, 2007). To determine the impact of SATB1-controlled gene signatures on Treg suppressive function, *SATB1* KO Treg cells as well as *AAVS1* and *FOXP3* KO control Treg cells were challenged in a T cell suppression assay (Fig. 5A). As expected, the ablation of FOXP3 had the strongest effect on Treg suppressive capacity. However, in all cell ratios tested, *SATB1* KO Treg cells also had a significantly reduced suppressive capacity compared to AAVS1 control edited cells (Fig. 5B, C). The KO Treg cell numbers were not negatively affected in these assays, excluding a reduced inhibition of Tconv cell proliferation based on diminished Treg survival and/or proliferation (Fig. 5D). Overall, human, *ex vivo* expanded *SATB1* KO Treg cells are phenotypically and functionally compromised.

**Figure 5.**
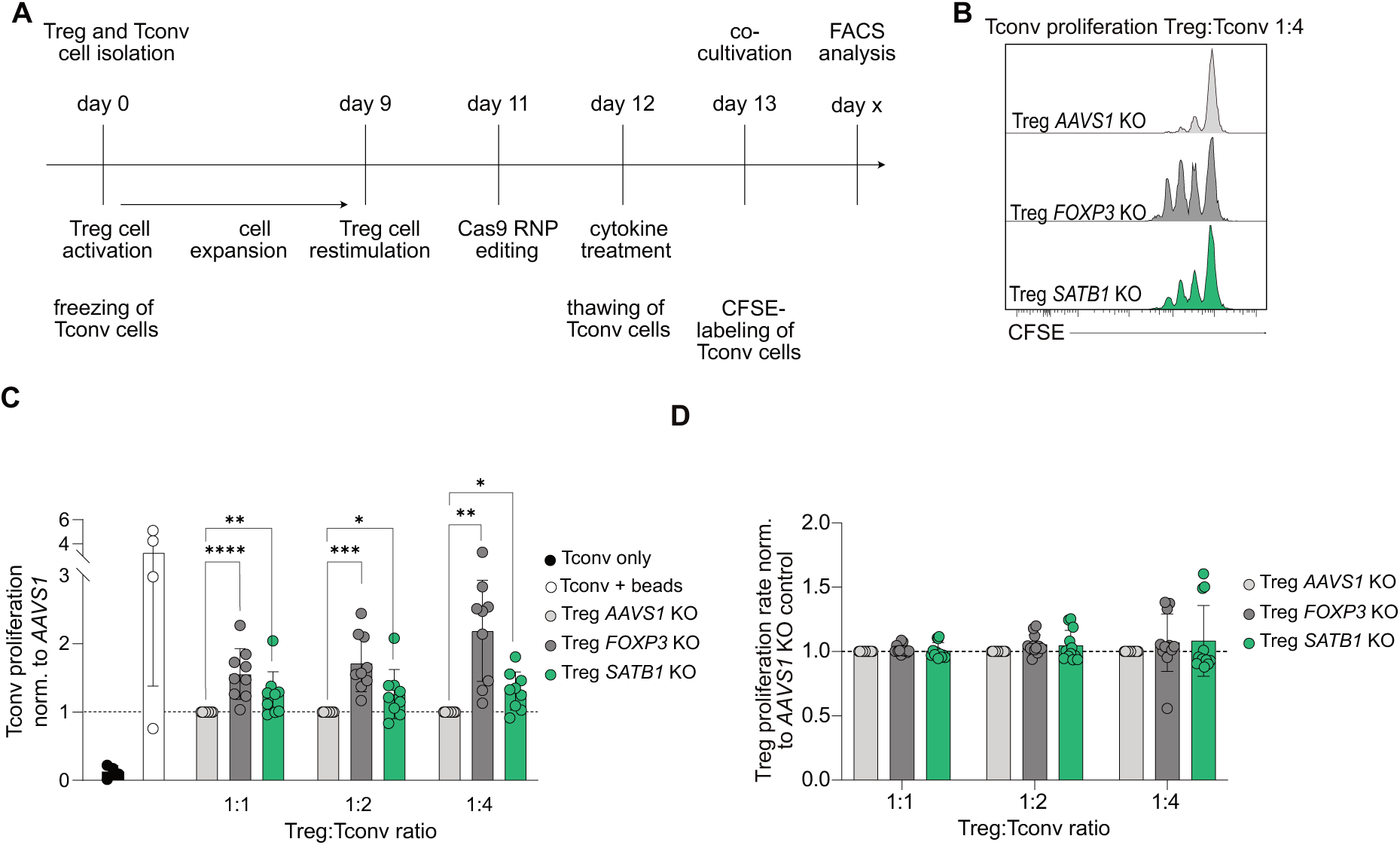
*SATB1* KO impairs Treg suppressive function. **(A)** Workflow of Treg suppression assay. Treg cells were isolated, expanded and CRISPR edited. Tconv cells were frozen directly after isolation and thawn on day 12 and rested overnight. Rested Tconv cells were labeled with CFSE and cultivated with KO Treg cells in different Treg:Tconv cell ratios (1:1, 1:2 and 1:4). Final flow cytometry quantification of CFSE-dilution was performed when 40-60 % of Tconv cells had undergone at least one cell division. **(B)** Representative histograms of Tconv cell proliferation cultivated with *AAVS1*, *FOXP3* or *SATB1* KO Treg cells in a Treg: Tconv cell ratios of 1:4. **(C)** Bar plots indicate mean with SD of Tconv cell proliferation rate normalized to *AAVS1* KO control condition. n = 3, with technical triplicates, two-way ANOVA with Tukey’s multiple comparisons test, * p<0.05, ** p<0.01, *** p<0.001, **** p<0.0001. **(D)** Bar plots indicating mean with SD of Treg proliferation rate normalized to *AAVS1* KO control condition. n = 4, with technical triplicates.

### *SATB1* KO in CD4 Tconv cells increases anti-tumor efficacy of CAR T cell products

Flow cytometry and integration of RNA-seq/ATAC-seq data revealed an enhanced T cell activation profile, as well as higher STAT5B levels after *SATB1* ablation. These changes could hint towards increased cell survival and expansion of Tconv cells after SATB1 ablation, which could potentially be beneficial for CAR T cell therapies.

To evaluate the effector functions of *SATB1* KO Tconv cells, we equipped them with anti-CD19 CAR receptors and challenged them in *in vitro* killing assays. *AAVS1* and *SATB1* KO CD4 CAR Tconv cells were cultivated with CD19-expressing Nalm6-FFLuc-GFP tumor cells in different cell ratios. *SATB1* ablation did not negatively affect the killing capacities of these cells (Fig. 6A). Interestingly, we observed an enhanced expansion of *SATB1* KO CAR Tconv cells in this assay in line with the previously identified gene signature indicating positive regulation of T cell expansion (Fig. 6B).

**Figure 6.**
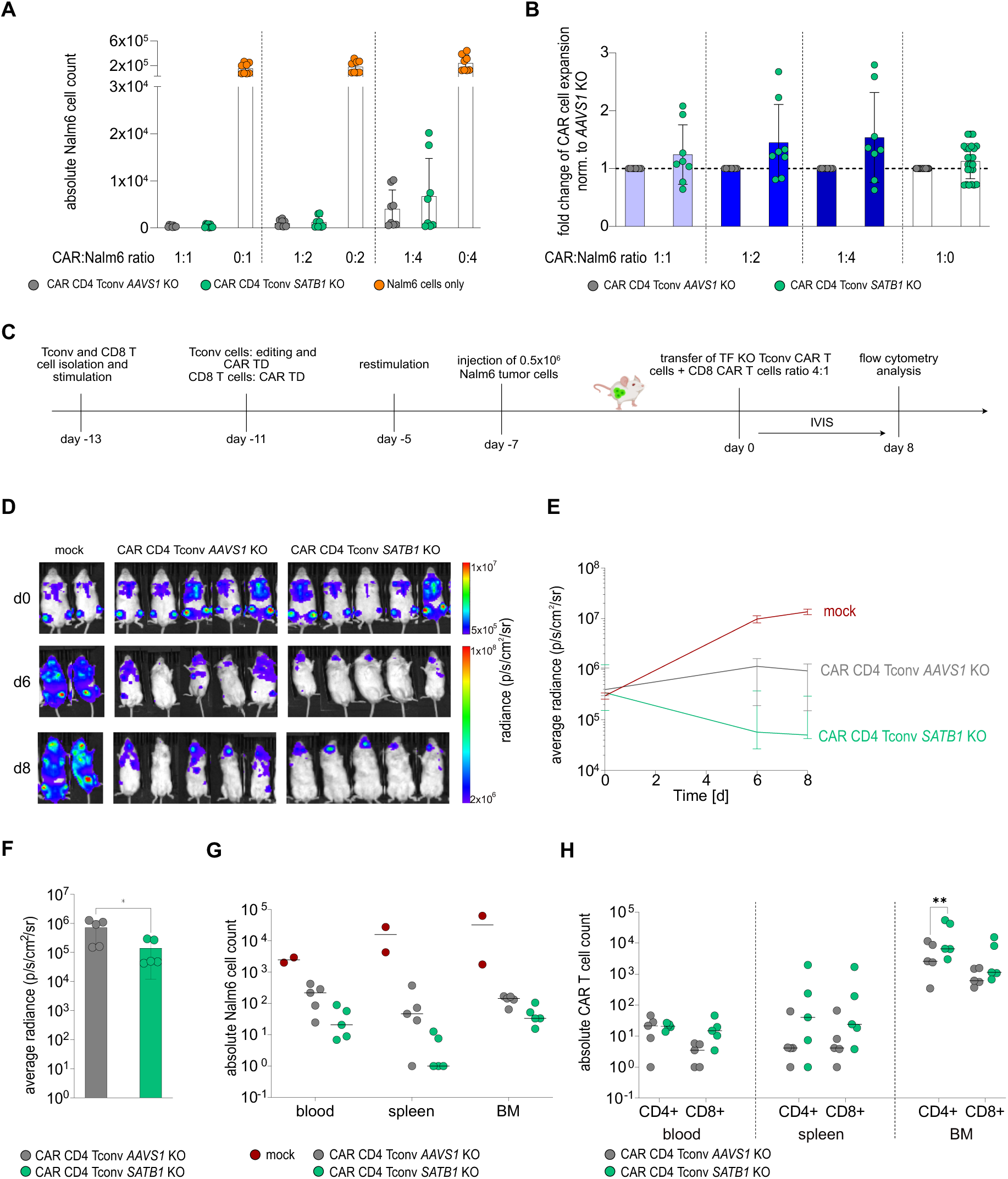
*SATB1* KO enhances CAR Tconv cell-mediated tumor clearance. **(A, B)** *AAVS1* and *SATB1* KO Tconv cells were retrovirally transduced with antiCD19-CAR and cocultured with or without human CD19+ Nalm6-FFLuc-GFP tumor cells in different CAR T cell to tumor cell ratios (1:1, 1:2 and 1:4). Tumor and CAR T cell counts were analyzed after 72 h of co-culture. **(A)** Bar plot of absolute counts of Nalm6-FFLuc-GFP tumor cells co-cultured with or without *SATB1* or *AAVS1* KO CAR Tconv cells. n = 6 with technical replicates, bar graphs indicate mean values with SD. **(B)** Bar plot indicates fold changes of CAR T cell expansion. Means with SD are shown. Expansion of *SATB1* KO CAR Tconv cells normalized to *AAVS1* KO control CAR Tconv cells. n = 6 with technical replicates, one-way ANOVA with Tukey’s multiple comparisons test, * p<0.05, ** p<0.01, *** p<0.001, **** p<0.0001. **(C)** Schematic workflow of *in vivo* functional validation of *SATB1* KO CAR CD4 Tconv cells. Nalm6-FFLuc-GFP tumor cells were injected into NSGS mice. After one week, *AAVS1* or *SATB1* KO Tconv cells transduced with anti-CD19-CAR retrovirus were adoptively co-transferred with anti-CD19-CAR-transduced CD8 T cells in a CD4 KO:CD8 cell ratio of 4:1 into Nalm6-FFLuc-GFP tumor-bearing NSGS mice. Tconv and CD8 T cells without CAR transduction served as mock control. Bioluminescence imaging (IVIS) was performed over time to monitor tumor development. Nalm6-FFLuc-GFP and CAR T cells were quantified in blood, spleen and bone marrow at day 8. **(D)** Bioluminescence images of Nalm6-FFLuc-GFP-bearing mice treated with mock, *AAVS1* KO, or *SATB1* KO CAR CD4 Tconv cells co-transferred with CAR CD8 T cells. **(E)** Average radiance over time. Tumor burden was quantified as the maximum photon per second per cm^2^ per steradian without unspecific head signal. Median with SD of average bioluminescence signal on day 0, 6 and 8 is indicated. n = 2-5. **(F)** Bar plot of average radiance. Tumor burden was quantified as the maximum photon per second per cm^2^ per steradian by excluding unspecific head signals. Mean with SD of average bioluminescence signal on the endpoint is indicated. n = 5, bar graphs indicate mean values, unpaired t-test, * p<0.05. **(G)** Absolute Nalm6-FFLuc-GFP tumor cell count measured via flow cytometry analysis of 100 µl blood, spleen and bone marrow (BM), n = 2-5, median is indicated. **(H)** Bar plots indicating median of absolute CAR T cell count of KO CD4 and CD8 T cells in 100µl blood, spleen and bone marrow (BM). n = 5, two-way ANOVA with Šídák’s multiple comparisons test, * p<0.05, ** p<0.01.

Next, we challenged these cells *in vivo*. We based this experimental outline on the publication by Ding and colleagues that expressed a constitutively active form of STAT5 specifically in anti-CD19 CAR CD4 Tconv cells. These so-called *CASTAT5* CAR T cells underwent robust expansion and cleared tumors efficiently (Ding *et al*, 2020). Shortly, *AAVS1* or *SATB1* KO CD4 T cells were transduced with a CD19-targeting CAR and injected together with unedited CD8 CAR T cells into Nalm6-FFLuc-GFP tumor-bearing NSGS mice. SATB1 has been described to prevent premature CD8 T cell exhaustion (Stephen *et al*, 2017). For that reason, CAR CD8 T cells were applied unedited to avoid biased results in tumor growth. 6 and 8 days after adoptive transfer, CAR T cell levels and tumor growth were analyzed (Fig. 6c). Interestingly, tumor clearance was more efficient in mice injected with *SATB1* KO CD4 CAR Tconv cells compared to *AAVS1* KO cells (Fig. 6D, E). The average radiance was significantly reduced at the endpoint on day 8 (Fig. 6F). In blood, spleen, and bone marrow a tendency toward fewer tumor cells could be observed in the *SATB1* KO condition (Fig. 6G). The counts of CD4 and CD8 CAR T cells in blood and spleen were similar in the *AAVS1* and *SATB1* KO conditions. However, in the bone marrow, the preferential site of Nalm6 tumor cell accumulation, an expansion of *SATB1* KO CD4 CAR T cells could be observed. CD8 CAR T cell numbers remained stable in the animals challenged with either *AAVS1* KO or *SATB1* KO CD4 CAR T cells (Fig. 6H). Our results show an improved functionality of CD4 CAR T cells after *SATB1* KO, which affirms the relevance of the CD4 T cell compartment for CAR therapy approaches.

## DISCUSSION

The KO or overexpression of transcription factors for improved CAR T cell function is currently explored in the context of T cell exhaustion, T cell activation, and modulating the plasticity of these cells, which includes overexpression approaches like constitutively activating STAT5 and c-JUN or ablation of *NR4A1* and *TOX* (Dai *et al*., 2024). Also, in CD4 T cells, especially Treg cells, transcription factors are targets of interest, namely the overexpression of FOXP3 and Helios for enhanced stability and suppressive function (Bittner *et al*, 2023). Here, we dissected the function of SATB1 in Treg and CD4 CAR T cell function. To be able to apply SATB1 purposefully to genetic engineering approaches, we asked whether SATB1 transcriptional signatures are subset-specific or largely conserved in CD4 T cells. In this study, we detected cell-specific SATB1 gene regulatory patterns in both Treg and Tconv, with a considerably larger set of affected genes in Treg cells. However, the SATB1-regulated pathways are mostly conserved in these two T cell subsets. KO of *SATB1* resulted in dysregulated JAK-STAT signaling and changes in cytokine/cytokine receptor interactions, which are closely intertwined.

We observed slightly reduced FOXP3 levels in Treg cells after SATB1 ablation; however, the direct comparison of *SATB1* KO to *FOXP3* KO Treg cells showed a clearly distinct gene regulatory pattern, which included, for example, increased levels of IL-10. Global analysis by ATAC-seq/RNA-seq highlighted dysregulation of multiple STATs in *SATB1* KO Treg cells. STAT3, STAT5b, and STAT6 were upregulated, whereas STAT1 levels were reduced. All of these proteins have been discussed to different extents in murine Treg cell function. Stat3 regulates the balance between Treg and Th17 cells by suppressing Th17 cell differentiation and negatively regulating Foxp3 expression (Qin *et al*, 2024). In multiple sclerosis, Stat3 inhibition can restore the balance between Treg and Tconv cells (Aqel *et al*., 2021). Stat6 signaling has been described as a negative regulator of Treg cell induction, stability, and suppressive function (Arroyo-Olarte *et al*, 2023). The observed upregulation of STAT5b and reduced levels of STAT1 can potentially counteract these destabilizing effects mediated by STAT3 and STAT6. Stat5 is a positive regulator of Foxp3 expression (Yao *et al*, 2007), whereas murine *Stat1* KO Treg cells have increased suppressive function *in vitro* and *in vivo* (Ma *et al*, 2011). Overall, changes in JAK-STAT signaling potentially contributed to an altered cytokine profile shifted towards pro-inflammatory cytokines.

In contrast to Treg cells, *SATB1* KO Tconv cells displayed reduced levels of pro-inflammatory cytokines. Several publications describe SATB1-mediated regulation of “cytokines cytokine receptor interactions” in different experimental contexts. Previously, SATB1 has been characterized as a driver of Th2 cell induction (Ahlfors *et al*., 2010; Cai *et al*, 2006). In our data, the picture is less clear. *SATB1* KO in Tconv resulted in a downregulation of GATA3 and IL-4, but also in an upregulation of classical Th2 cytokines including *IL5* and *IL9* in Tconv cells. However, the *in vitro* culture conditions applied here – strong stimulation by antiCD3/CD28 and high doses of IL-2 – favor Th0/Th1 cell phenotypes. Gupta et al. described murine *Satb1* KO Tconv cells to be more susceptible to Treg cell suppression *in vitro* and *in vivo* due to lower expression levels of CD25 and IL-2, while there was no effect on T cell proliferation (Gupta *et al*., 2022). In our experimental setting with human Tconv cells, we also detected decreased IL-2 secretion with only slight effects on IL-2 receptor expression. However, integration of RNA-seq and ATAC-seq data of *SATB1* KO Tconv highlighted a heightened activation profile. This could be confirmed on a functional level by increased *in vitro* and *in vivo* expansion of CD4 CAR T cells. A heightened activation profile could be (partially) driven by STAT5B. Constitutively active STAT5 has been shown to boost CD4 CAR functionality by increasing cell survival and expansion (Ding *et al*., 2020). In our experiments, *SATB1* KO CD4 CAR Tconv cells outperformed *AAVS1* KO control cells in a Nalm6 tumor model.

Together Treg destabilization and simultaneous increased activation of CD4 CAR T cells could be an interesting strategy to boost the efficiency of adoptive CAR T cell therapies. However, additional work is necessary to fully assess the long-term effects and to ensure safety.

## Acknowledgments

We thank the members of the Schumann laboratory for helpful suggestions and technical assistance. We thank Jeffrey A. Bluestone for critical feedback on our study. We especially thank Immanuel Andrä, Katharina Hofmann, Tanja Roßmann-Bloeck and Corinne Angerpointner for cell sorting assistance. We acknowledge support by the Core Facility “CyTUM MIH” in Cell Sorting and are thankful for the participation of the DFG in instrument funding within Project Nr. 659391 and 659389. We kindly thank Stanley Riddell for the Nalm6-FFLuc-GFP cell line. This work was in part funded by the Intramural Research Program (IRP) of the National Institute of Diabetes and Digestive and Kidney Diseases (M.H.). This work was funded by the Deutsche Forschungsgemeinschaft (DFG, German Research Foundation) grants SFB1054/3 - 210592381 and SFB-TRR 338/1 2021-452881907 to K.S., D.K. and D.H.B. and SFB-TRR 355/1 2022 – 490846870 to I.S., C. D. and K.S.

## Author contributions

S.K. and K.S. designed research; S.K., L.D., and J.R., performed research; K.S., S.K. and C.S. designed ATAC-seq experiment; K.S., S.K. designed RNA-seq experiment; S.K. performed ATAC-seq and RNA-seq experiments; S.K., E.D.L. M.H., M.L. C.S. analyzed ATAC-seq data; S.K., E.D.L. M.H. analyzed RNA-seq data; M.W. provided cell material; D.K. provided critical feedback to data interpretation; I.S. and C.D. supported functional Treg cell analysis, L.W., S.K., K.S. and D.H.B. designed and analyzed *in vivo* CAR T cell experiment; L.W. and S.K. performed *in vivo* CAR T cell experiment; K.S. wrote the manuscript with the support of all authors.

## MATERIAL & METHODS

### Mouse model

NSGS mice (NOD.Cg-Prkdcscid Il2rgtm1Wjl Tg (CMV-IL3,CSF2,KITLG)1Eav/MloySzJ) (female, 6-8 weeks old, 18 – 22 g) were acquired from The Jackson Laboratory and kept at the mouse facility at the Technical University Munich, Institute for Medical Microbiology, Immunology and Hygiene. The mice were housed in groups under special, pathogen-free conditions at a constant temperature of 20 °C with constant availability of food and water and subjected to a 12:12 day/night cycle. Littermates of the same sex were randomly allocated to the experimental groups. The performed animal experiments were approved by the district government of Upper Bavaria (Department 5—Environment, Health and Consumer Protection ROB-55.2-2532.Vet_02-18-162).

### Primary human T cells

Buffy coats were collected by the Bavarian Red Cross, Donas GmbH or the German Heart Center Munich, with the approval of the local institutional review board (Ethics Committee TUM School of Medicine, Technical University of Munich) and with the informed consent of the patients. Information about age and gender of donors is not available. The study conforms to the standards of the Declaration of Helsinki. PBMCs were isolated using gradient density centrifugation with Pancoll (PAN-Biotech) and cultured in cRPMI as described in detail below.

### Isolation and expansion of primary human T cells

PBMCs were isolated with Pancoll (Density: 1.077 g/ml; PAN-Biotech) and SepMate^TM^ tubes (STEMCELL Technologies) out of buffy coats. CD4 T cells were pre-enriched with MojoSort^TM^ Human CD4 T cell Isolation kit (BioLegend). For Treg and Tconv cell isolation, CD4 cells were flow cytometry-sorted (anti-human CD4-Pacific Blue™ (clone SK3, BioLegend), anti-human CD25-APC (clone BC96, BioLegend), and anti-human-PE CD127 (clone A019D5, BioLegend)) based on CD4 expression and CD25^high^CD127^low^ (Treg cells) or CD25^low^CD127^high^ (Tconv cells) using a FACS Aria III (Software: FACS Diva 8.0; Becton Dickinson) or a MoFlo Astrios EQ cell sorter (Software: Summit 6.3; Beckman Coulter).

Isolated Treg and Tconv cells were stimulated with Dynabeads™ Human T-Activator CD3/CD28 (Gibco, 25 µl per 1×10^6^ cells) in a cell:bead ratio of 1:1 and cultivated in complete Roswell Park Memorial Institute medium (T cell medium (TCM), consisting of RPMI 1640 (Thermo Fisher Scientific) supplemented with 5 mmol/l HEPES (PAN-Biotech), 2 mmol/l glutamine (PAN-Biotech), 50 µg/ml penicillin/streptomycin (PAN-Biotech), 5 mmol/l nonessential amino acids (PAN-Biotech), 5 mmol/l sodium pyruvate (PAN-Biotech) and 10% FCS (fetal calf serum, Gibco) at 37°C. Treg cells were cultured at 0.25×10^6^ cells/ml with 600 U/ml IL-2 (Peprotech) and Tconv cells at 0.5×10^6^ cells/ml with 200 U/ml IL-2 (Peprotech).

### CRISPR/Cas9 RNP editing

On day 9 post-isolation, 0.7×10^6^ Treg and Tconv cells were stimulated with plate-coated anti-human CD3 (BioLegend, clone UCHT1, 10 µg/ml in 150 µl PBS/48-Well) and soluble anti-human CD28 (BioLegend, clone CD28.2, 5 μg/ml) in the presence of 600 or 200 U/ml IL-2 (Peprotech). 48 h after activation, 0.3×10^6^ to 1×10^6^ cells were nucleofected with 4 µl Cas9 RNPs and 1 µl 100 µM electroporation enhancer (Sigma-Aldrich) in 20 µl buffer P3 with supplement (Lonza). Cas9 RNPs were generated by mixing 100 µM crRNA (IDT, protospacer sequence: Table S1) and 100 µM tracrRNA (IDT) in a 1:1 ratio and incubated for 5 min at 96°C. After cooling down to room temperature (RT) 40 μM *Streptoccocus pyogenes* Cas9-NLS (in-house production; Cas9 expression plasmid pMJ915 (Addgene), (Lingeman *et al*, 2017)) was slowly added to the 50 μM crRNA:tracrRNA duplexes and incubated for 15 min at RT. Nucleofections were performed with Amaxa 4D-Nucleofector (Lonza) using program EH-115 for Treg cells and DK-100 for Tconv cells. After nucleofection, cells received additional stimulation using 25 µl ImmunoCult^TM^ (STEMCELL Technologies)/1×10^6^ cells and 600 or 200 U/ml IL-2. Treg cells were cultivated with TCM whereas Tconv cells were cultured in X-VIVO medium (X-VIVO 15 Lonza, supplemented with 50 µg/ml penicillin/streptomycin (PAN-Biotech) and 10% FCS (Gibco)). One day after nucleofection, expanded Treg and Tconv cells were split into two separate conditions treated either with 600 or 200 U/ml IL-2, respectively, or with additionally 1 µg/ml hIL-12 (Miltenyi).

### Flow cytometry analysis

Cells were stimulated with 6.25 ng/ml PMA (Sigma-Aldrich), 1 µg/ml Ionomycine (Sigma-Aldrich) and 1:1200 GolgiPlug™ (BD Bioscience) for 5 h in 160 µl TCM. Surface antibody staining was performed in 30 µl PBS with respective amounts of fluorophore-labeled antibodies at 4 °C in the dark. FOXP3 Fix/Perm Buffer Set (BioLegend) was used to fixate cells for 30 min at RT in the dark. Intracellular staining was performed in 30 µl PERM buffer (1:10 diluted with PBS) for 30 min at RT in the dark. Phenotyping of KO Treg and Tconv cells was performed with Zombie NIR™ (BioLegend), anti-human CTLA-4-APC/Fire750 (clone L3D10MQ1, BioLegend), anti-human IL-2-BV650 (clone 17H12, BioLegend), anti-human IL-10-PE (clone JES-9D7, BioLegend), anti-human FOXP3-AF488 (clone 206D, BioLegend), anti-human IFNγ-BV785 (clone 4S.B3, BioLegend), and anti-human Helios-PE-Cy7 (clone 22f6, BioLegend). Cell acquisition was conducted on a Cytoflex LX or Cytoflex S instrument (software CytExpert 2.4, Beckman Coulter) or Aurora analyzer (software SpectroFlo®, Cytek). Cells were analyzed with FlowJo (v10.6). Optionally, FCS files were analyzed in Omiq (OMIQ) and the respective compensation matrix of the FlowJo analysis was applied. Cells were pregated for viable singlets and the data sets subsetted dependent on the respective CD4 T cell subset and cytokine treatment. 4000 viable singlets of each sample were included in the analysis. opt-SNE clustering algorithm embedded in the software was applied for optimized local structure resolution based on all FACS markers (CD25, CTLA-4, TIGIT, FOXP3, Helios, IL-10, IL-2, IL-4, IFN-γ) (Belkina *et al*, 2019).

### Legendplex Assay

To determine the absolute amount of secreted cytokines LEGENDplex^TM^ HU Essential Immune Response Kit (Biolegend) containing capture beads for human IL-4, IL-2, TNF-α, IL-17A, IL-6, IL-10, IFNγ and active TGFβ was applied. Supernatant of the samples and standards were measured in duplicates. 12.5 µl of the prepared standard dilutions or samples and 12.5 µl assay buffer were transferred in a v-bottom plate. 12.5 µl of the diluted capture bead mixture were added and incubated over night at 4 °C in the dark. After three washing steps, 12.5 µl of the detection antibodies were added and the samples incubated for 1 h in the dark, while shaking at 800 rpm at RT. Without washing 12.5 µl of the Streptavidin-PE solution were added and incubated for another 30 min at RT while shaking at 800 rpm. For acquisition, the plate was washed and samples were resuspended in 150 µl of 1X wash buffer. Samples were acquired at the Cytoflex LX or Cytoflex S instrument (software CytExpert 2.4, Beckman Coulter) that was previously set up with the Setup Beads 3 (Raw beads) and Setup Beads 2 (PE beads), provided by the manufacturer.

### Amplicon Sanger Sequencing

To isolate the genomic DNA out of the genetically modified cells, 10 µl cell suspension containing approximately 1×10^4^ cells was added to 30 µl QuickExtract DNA Extraction Solution (Biosearch Technologies) and incubated for 6 min at 65°C followed by 2 min at 96°C. One PCR reaction contained 12.5 µl of GoTaq® Long PCR Master Mix (Promega), 1.25 µl of 10 µM forward and reverse primer, respectively (Sigma-Aldrich), 2 µl of extracted DNA solution and 8 µl of H2O. The primers were designed to generate an amplicon of approximately 750 bp around the expected gRNA cut site (see Table S2). The thermocycler setup was as follows: initial denaturation at 95 °C, 3 min; 14 cycles of 98°C 20 s, 65°C 20s, 72°C 60 s with a touchdown of -1 C/4s and 29 cycles of 98°C 20 s, 58°C 20 s, 72°C 60 s. Sanger sequencing was performed by Microsynth AG and KO efficiencies were determined using the TIDE webtool (Brinkman *et al*, 2014).

### Amplicon NGS

The indel patterns of CRISPR/Cas9-edited human T cells were determined by amplicon PCR followed by NGS. Primers were designed to result in an amplicon length of 350 to 500 bps using Benchling (see Table S3). A 25 µl PCR reaction was performed as described above with the respective primers. The PCR products were cleaned up using AMPure beads according to the manufacturer’s recommendations (Beckmann Coulter) and eluted in 52.5 µl 10 mM Tris. For barcoding, 2 µl of the purified DNA were added to 10 µl of 2x GoTaq Long PCR Master Mix, 2 µl of Nextera XT index 1 (i7) primer, 2 µl of Nextera XT index 2 (i5) primer (Illumina) and 4 µl of H2O. The PCR reactions were heated up to 95 °C for 3 min, followed by 8 cycles of 95°C for 30 s, 55°C for 30 s and 72°C for 30 s and an elongation step for 5 min at 72°C. The barcoded PCR products were again purified with AMPure beads and eluted in 27.5 µl 10 mM Tris. Cleaned-up PCR products were quantified using the SpectraMax Quant AccuBlue HighRange dsDNA Kit (Molecular Devices) on the SpectraMax 3x instrument (Molecular Devices). Equal amounts of DNA/sample were pooled and sequenced on an Illumina MiSeq instrument (Illumina) with a MiSeq Reagent Nano Kit v2 (500-cycles) (Illumina) according to the manufacturer’s recommendations. NGS sequencing results were analyzed using CRISPResso (version 2.2.7) with the following prompt: CRISPRessoBatch –batch_settings [name.batch] –amplicon_seq [sequence amplicon] -g [sequence gRNA] -n nhej -gn [name gRNA] -w 30 –skip_failed -o [name of output folder].

### RT-qPCR

0.5 to 1×10^6^ viable KO Treg and Tconv cells treated with IL-12 were flow cytometry-sorted on day 4 post-nucleofection. RNA isolation was performed with the Quick-RNA Microprep Kit (Zymo Research). cDNA was generated out of 500 ng to 1 mg of RNA. RNA was incubated at 70 °C for 5 min with 150 ng random hexamer primers (Promega). Nuclease-free water was added to the RNA-primer mix to a final volume of 18 µl and mixed with the RT reaction mix: 1 µl reverse transcriptase (M-MLV, Promega), 5 µl M-MLV RT buffer (Promega) and 1 µl or 10 mM dNTPs (New England Biolabs). cDNA synthesis was performed with the initial primer extension step at 22 °C for 10 min, followed by the reverse transcriptase reaction step at 50 °C for 50 min, and finalized with the heat inactivation at 70 °C for 15 min. cDNA was diluted 1:10 to quantify FOXP3 and SATB1 expression and 1:10000 for the 18S rRNA used as a housekeeping gene. 5 µl 2x GoTaq qPCR master mix (Promega) was mixed with 0.5 µl of forward and reverse primer (see Table S4) 10 µM primer stock and 4 µl of the diluted cDNA. The qPCR protocol was as follows: 95 °C for 300 s, 40 cycles of 95 °C for 10 s, and 60 °C for 30 s on CFX Connect Real-Time PCR Detection System (Biorad). Data analysis of the RT-qPCR was done by the ΔΔCT method. After normalization to a housekeeping gene, the cycle threshold (CT) values were compared with a sample control.

### RNA-seq

0.5 to 2×10^6^ viable Treg or Tconv cells were flow cytometry-sorted by applying propidium iodide (BioLegend) 1:500 to the cells. RNA was isolated with the Quick-RNA Microprep Kit (Zymo Research). Novogene, UK, performed library preparation, sequencing, and analysis. In brief, library preparation was performed as follows: mRNA was purified from total RNA using poly-T oligo-attached magnetic beads. After fragmentation, the first strand cDNA was synthesized using random hexamer primers, followed by the second strand cDNA synthesis using dTTP for the non-directional library (Parkhomchuk *et al*, 2009). The non-directional library was ready after end repair, A-tailing, adapter ligation, size selection, amplification, and purification.

Qubit and real-time PCR were used for quality control and quantification, and bioanalyzer measurements were applied for size distribution detection. Quantified libraries were pooled and sequenced using a Novaseq 6000 X Plus pair-end 150 sequencing strategy.

Raw data (raw reads) of fastq format were cleaned by removing reads containing adapter, poly-N, and low-quality reads from raw data. The alignment to the reference genome was performed with Hisat2 v2.0.5 (Mortazavi *et al*, 2008). FeatureCounts v1.5.0-p3 (Liao *et al*, 2014) was used to count the reads numbers mapped to each gene. Then, the FPKM of each gene was calculated based on the length of the gene and the read count mapped to this gene. Read counts were normalized and transformed using VST with DESeq2 (v1.44.0). (Love *et al*, 2014) Two additional targets (“H” and “D”) were incorporated into the batch correction process to enhance the robustness of the dataset. However, their individual results were excluded from the scope of this study. The transformed data were visualized using PCA and plotted with the R package ggplot2 (v3.5.1) (Wickham, 2010).

Differential expression analysis was analyzed with DESeq2 (v.1.20.0) (Anders & Huber, 2010; Love *et al*., 2014). The resulting p-values were adjusted using Benjamini and Hochberg’s approach to control the false discovery rate. Differentially expressed genes (DEGs) were defined as genes with FDR < 0.05 and log2 fold change > 0.5. To visualize the DEGs, volcano plots were drawn using the Python package ngs-toolkit (v0.25.1) (https://github.com/afrendeiro/toolkit, André F. Rendeiro).

### ATAC-seq

At day 4 after Cas9 RNP nucleofection, 5.5×10^4^ flow cytometry-sorted, live *AAVS1* or *SATB1* KO Treg and Tconv cells treated with IL-12 were washed with 500 µl ice-cold ATAC buffer (10 mM Tris-HCl pH 7.4; 10 mM NaCl in Ambion H2O). Cell lysis was performed with 50 µl cold ATAC buffer supplemented with 10 % NP40 (Sigma-Aldrich), 10 % TWEEN-20 (Sigma-Aldrich), and 1 % Digitonin (Promega) for 3 min at 4 °C. 1 ml cold ATAC buffer with 0.1 % TWEEN-20 (Sigma-Aldrich) was used to stop the reaction. The cells were resuspended in 50 µl Transposase Mastermix: TD buffer (Illumina), 16.5 µl DPBS, no calcium, no magnesium (Thermo Fisher Scientific), 0.5 µl 1 % Digitonin (Promega), 0.5µl 10 % Tween-20 (Sigma-Aldrich) and 100 nM Transposase (Illumina) and incubated for 30 min at 37 °C. The cleanup of the transposed fragments was performed with the Zymo DNA Clean and Concentrator-5 Kit (Zymo Research) according to the manufacturer’s instructions. 23 µl DNA Elution Buffer (Zymo Research) were added to elute the transposed fragments from the columns. To determine the optimal number of PCR cycles, 10 % of the purified ATAC-seq DNA sample was subjected to qPCR to avoid over-amplification of libraries.

5 µl of NEBNext High Fidelity 2x PCR Mastermix (NEB), 1.9 µl nuclease-free water (Invitrogen), 0.5 µl of 25 µM Ad1.1 primer and Ad2.1 primer (Corces *et al*, 2017) and 0.1 µl of 100 x SYBR green (Thermo Fisher Scientific) was mixed with 2 µl of the purified sample. The qPCR protocol was as follows: 5 min at 72 °C followed by 30 sec at 98 °C and 25 cycles of 10 sec at 98 °C, 30 sec at 63 °C and 1 min at 72 °C. The optimal cycle number for the final enrichment PCR was determined by rounding up the Ct value of each sample individually. For the final enrichment PCR, 20 µl of the purified sample was mixed with 25 µl NEBNext High Fidelity 2x PCR Mastermix (NEB), 2.5 µl of 25 µM Ad1.xx primer and 2.5 µl of 25 µM Ad2.xx primer using different barcode combinations for each sample (see Table S5). The PCR program was performed with the individual cycling numbers described above, and adding a final extension step for 1 min at 72 °C. The PCR product was purified with the Zymo DNA Clean and Concentrator-5 Kit after the vendor’s manual using 15 µl DNA Elution Buffer.

The size selection of 150 to 580 bp fragments was performed with AMPure XP Beads (Beckman Coulter). 0.47 x AMPure XP Beads were mixed with the purified tagmented sample and incubated for 10 min at RT. The sample was placed on a magnet for 5 min, and the supernatant was transferred into a new collection tube to eliminate smaller fragments. To remove larger fragments, a final concentration of 1.8 x vortexed AMPure XP beads were mixed with the supernatant and incubated 10 min at RT. After 5 min incubation time on the magnet, the supernatant was discarded, and the beads coupled to the DNA were washed 2 times with 180 µl 80 % ethanol. The beads were air-dried for 4 min, and the DNA was eluted using 15 µl TRIS pH 8 (5 min, RT). To remove the beads, the sample was placed for 5 min on the magnet, and the supernatant was transferred to a new collection tube.

The DNA concentration was measured with the Qubit dsDNA HS Assay Kit (Invitrogen). Samples were sent for sequencing to Novogene, UK. The raw data from ATAC-seq was analyzed as described before (Delacher *et al*, 2021).

### Preprocessing and analysis of ATAC-seq data

ATAC-seq reads were trimmed using Skewer (Jiang *et al*, 2014) and aligned to the hg19 assembly of the human genome using Bowtie2 (Langmead & Salzberg, 2012) with the ‘-very-sensitive’ parameter and a maximum fragment length of 2000 bp. Duplicate and unpaired reads were removed using the sambamba_v0.7. (Tarasov et al., 2015) ‘markdup’ command, and reads with mapping quality >30 and alignment to the nuclear genome were kept. All downstream analyses were performed on these filtered reads. For visualization purposes only, coverage files from filtered bam files were produced using deeptools_v3.5.1 (Ramirez *et al*, 2014) with the parameters ‘--binSize 10 --normalizeUsing RPGC --effectiveGenomeSize 3300000000 --extendReads 175’.

Peak calling for each sample was performed using MACS2 (Zhang *et al*, 2008) with the parameters ‘--nomodel --extsize 147’. Peaks overlapping blacklisted features as defined by the ENCODE project (The ENCODE Project Consortium 2012) were discarded. For the analysis of sample sets, a consensus region was created by merging the called peaks from all involved samples, and we quantified the accessibility of each area in each sample by counting the number of reads from the filtered bam file that overlapped each region.

DESeq2 was used on the raw count values for each sample and regulatory element to identify differential chromatin accessibility between samples after normalization of a matrix using the Variance Stabilization Transformation (VST) method and considering the donor as a covariate to remove batch effects. (Love *et al*., 2014) Two additional targets (“H” and “D”) were incorporated into the batch correction process to enhance the robustness of the dataset. However, their individual results were excluded from the scope of this study. Significant regions were defined as having an FDR-corrected p-value below 0.05 and absolute log2 fold change above 2. Peaks were assigned to their nearest transcription start site using the HOMER promoter annotation (Heinz *et al*, 2010).

### Integration of the RNA-seq and ATAC-seq data

The gene-to-peak correlation for ATAC-seq and RNA-seq data was done according to Thakore et al. (Thakore *et al*., 2024) for groups Tconv *SATB1* KO vs Tconv *AAVS1* KO and Treg *SATB1* KO vs Treg *AAVS1* KO. For each differentially expressed gene, a window of ±10 kb around the TSS was defined. All differentially accessible ATAC peaks of the respective groups intersecting this gene window by at least one base were assigned to the gene. When multiple peaks overlapped the same gene window, the peaks and their log2 fold changes were aggregated by calculating the mean. RNA counts and ATAC peaks were transformed using VST with DESeq2 (Love *et al*., 2014). The z-scores of the gene counts and their assigned peaks, along with their respective log2 fold changes, were visualized using the R library ComplexHeatmap (Gu *et al*, 2016). The source code used for this analysis is found at https://github.com/daisybio/Tconv-treg-satb1-analysis.

### Treg cell suppression assay

Treg cells were isolated, expanded, and edited with Cas9 RNPs as described previously. One day after Treg CRISPR-editing, a mixture of Tconv cells of different donors (standardized “responder T cells”) was thawed and rested overnight. On the following day, Tconv cells were labeled with CFSE as follows: Up to 1×10^7^ Tconv cells were washed with PBS and resuspended in 1 ml PBS. CFSE (2.78 µg/µl, BioLegend) was diluted 1:2000 in PBS. 1 ml of diluted CFSE was added to 1 ml cell suspension and incubated for 5 min at RT in the dark. To stop the staining, 2 ml TCM was added and incubated again for 1 min at RT. The cells were washed once. 1×10^5^ CFSE labeled Tconv cells were cultured together with 1×10^4^ Dynabeads™ Human T-Activator CD3/CD28 (Gibco) and different amounts of edited Treg cells ranging from 1:1, 1:2 and 1:4 Tconv:Treg cell ratios. The final readout was conducted after 40-60 % of CFSE-labeled Tconv cells divided at least once upon activation (see Fig. S4). Cytokine secretion of Treg cells after co-culture was quantified via flow cytometry: live dead Zombie NIR (BioLegend), anti-human FOXP3-PE (clone 206D, BioLegend), anti-human IL-2 Brilliant Violet 510™ (clone MQ1-17H12, BioLegend), anti-human IFNγ Brilliant Violet 785™ (clone 4S.B3, BioLegend), anti-human IL-10 PE/Dazzle™ 594 (clone JES-9D7, BioLegend) and, anti-human IL-4 Brilliant Violet 421™ (clone MP4-25D2, BioLegend).

### Retrovirus production

1.2×10^6^ RD114 cells (human rhabdomyosarcoma cell line) were seeded in 3 ml DMEM (PAN-Biotech) supplemented with 10 % FCS (Gibco) and 1 % Penicillin-Streptomycin (PAN-Biotech) in 6-well culture plates. 18 µg plasmid DNA (JCAR021 in pMP72, a mutant of JCAR017 (clone: FMC63) from Juno Therapeutics – a Bristol Myers Squibb company) and 15 µl CaCl2 solution (3.31 M, Sigma-Aldrich) were mixed with H2O in 150 µl. The mixture was added dropwise while vortexing to 150 µl transfection buffer (1.6 g NaCl (Sigma-Aldrich), 74 mg KCl (Sigma-Aldrich), 50 mg H2PO4 (Sigma-Aldrich), 1 g HEPES (Sigma-Aldrich) in final 100 ml H2O pH 6.76). After 20 min at RT, the transfection reagent was added dropwise to RD114 cells. The medium was exchanged after 4 h to TCM. Retrovirus was harvested 48 h and 72 h post-transfection.

### Nalm6-tumor cell killing assay

CD4 Tconv cells were isolated via flow cytometry-sorting and activated with ImmunoCult^TM^ (5 µl/ 1×10^6^ cells, STEMCELL Technologies) in the presence of 200 U/ml IL-2. After 48 h, CRISPR-editing was performed as indicated above. After 2 h of resting, the cells were transduced with retrovirus. Non-treated tissue culture plates were coated with 0.06 µg/ml RetroNectin (Takara) in 300 µl PBS in a 24 well-plate. 900 µl CAR retrovirus supernatant was coated on the RetroNectin-coated wells by centrifugation (2 h at 3000 x g and 32 °C). 700 µl supernatant was replaced by CD4 Tconv cell suspension (0.5×10^6^ cells/well) supplemented with 200 U/ml IL-2 in the same volume. After one week of cultivation, cell counts were adjusted according to the transduction rate. 2×10^4^ CAR-positive *AAVS1* or *SATB1* KO CD4 Tconv cells were cultured together with CD19^+^ Nalm6-FFLuc-GFP tumor cells (acute lymphoblastic leukemia (ALL)) in different T cell to tumor cell ratios (1:1, 1:2, and 1:4). CAR CD4 T cell expansion and tumor cell numbers were determined 24 h and 72 h after co-culture. For the final flow cytometry-based readout following reagents were used: 7×10^3^ 123count eBeads™ (Thermo Fisher Scientific), live dead Zombie NIR (BioLegend), anti-human CD4-PE (clone SK3, BioLegend), anti-human EGFR-APC (clone AY13, BioLegend), and Streptavidin-eF480 (eBioscience^TM^).

### Nalm6-FFLuc-GFP tumor model

7 days prior to T cell transfer, 6 to 10-week-old NSGS mice were injected with 0.5×10^6^ CD19^+^ Nalm6-FFLuc-GFP cells. Tumor growth in mice was quantified by IVIS imaging one day prior to T cell transfer. Mice were injected intraperitoneally with 150 mg/kg XenoLight D-Luciferin Potassium Salt (PerkinElmer) dissolved in PBS. After 5 min, mice were anesthetized with 2.5 % isoflurane RAS-4 Rodent Anesthesia system (Perkin Elmer) and imaged in the IVIS Lumina Imaging System (PerkinElmer LAS). The analysis was performed by quantification of photons/sec/cm^2^/sr with Living Image 4.5 software (PerkinElmer).

KO CD4 Tconv cells as well as non-edited CD8+ T cells transduced with JCAR021, were adjusted according to their transduction rate to a final cell ratio of 1.2x 10^6^ transduced TF KO CD4+ Tconv cells and 0.3x 10^6^ transduced CD8+ T cells. Mock control mice received CD4 Tconv cells and CD8 T cells without CAR. Tumor growth was determined once per week and at the endpoint by IVIS bioluminescence imaging. 8 days after T cell transfer, mice were sacrificed and lymphocytes in blood, spleen, and bone marrow were harvested. 100 µl blood was added to 10 ml ACT buffer (0.17 M NH4Cl (Sigma-Aldrich), 0.3 M Tris-HCl (Sigma-Aldrich) pH 7.5) and incubated for 10 min at RT. The lysis was stopped by adding 4 ml of cold TCM. The step was repeated with 5 ml ACT buffer for 5 min after pelleting the cells by 5 min at 1500 rpm. Spleens were mashed through a 100 µm cell strainer and ACT lysis was carried out with 5 ml ATC buffer for 5 min. The bone marrow was isolated out of femur and tibia of both legs followed by lysis with 3 ml ACT buffer for 3 min. Isolated cells were characterized by flow cytometry: 1×10^4^ 123count eBeads™ (Thermo Fisher Scientific) per condition, anti-human CD19-PE/Dazzle594 (clone HIB19, BioLegend), anti-human CD45-PB (clone T29/33, Dako), anti-human CD4-PacificOrange (clone RPA-T4, eBioscience^TM^), anti-human CD8-APC/Fire™ 750 (clone SK1, BioLegend), Streptavidin-FITC (BioLegend), anti-human EGFR-PE (clone AY13, BioLegend), live/dead staining with propidium iodide (BioLegend).

## Supplemental information

Supplementary Tables (combined in one excel file):

Table S1: gRNA sequences

Table S2: Primer sequences for quantification of CRISPR editing efficiencies by Sanger sequencing

Table S3: Primer sequences for quantification of CRISPR editing efficiencies by amplicon NGS

Table S4: RT qPCR primer sequences

Table S5: Sequences of barcoded ATAC-seq primers

Table S6: Pathway analysis based on RNA-seq data of *SATB1* KO Treg and Tconv cells

**Figure S1.**
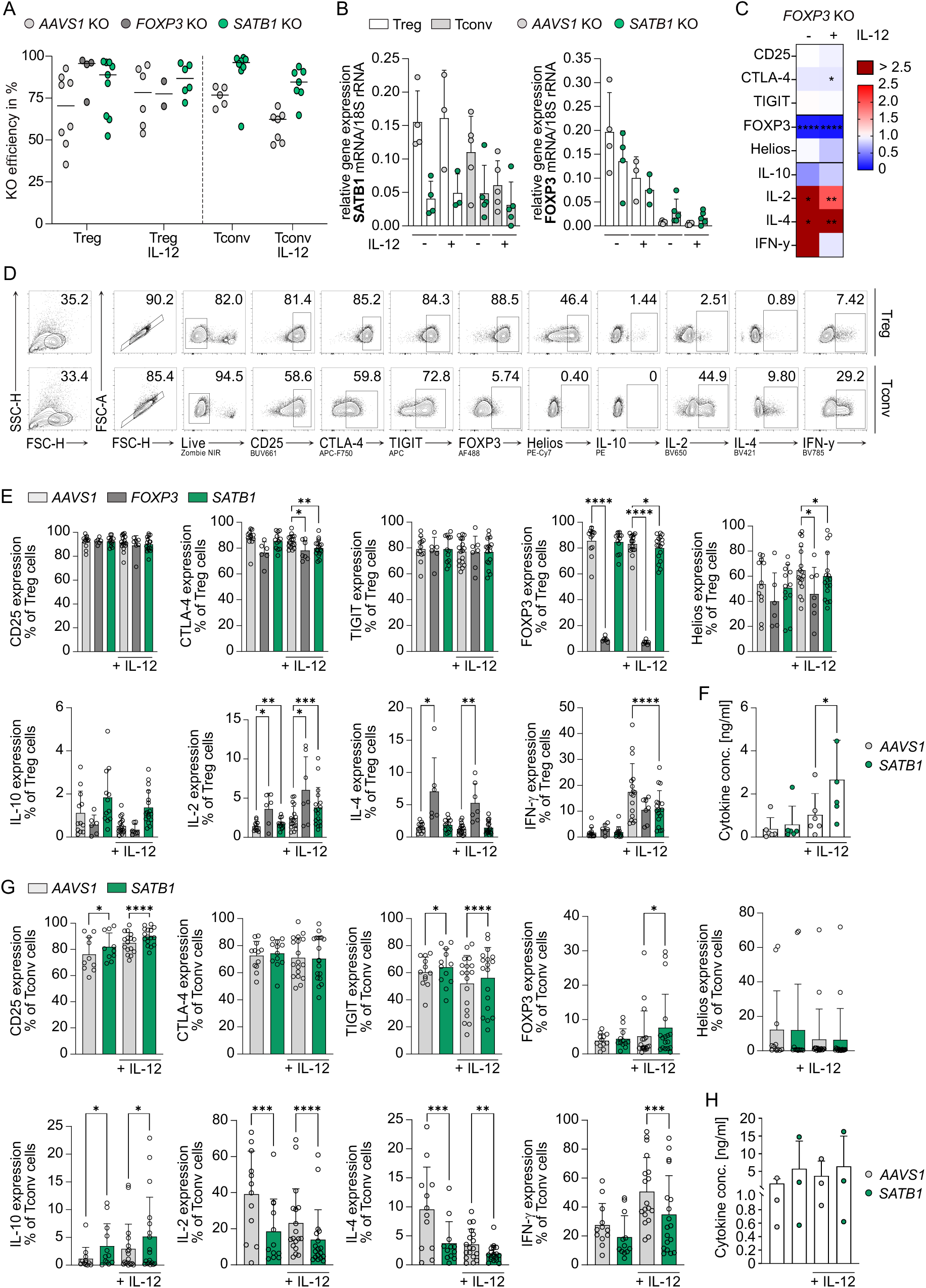
*SATB1* KO validation and overall phenotypic alterations in SATB1 ablated T cell subsets. Human Treg and Tconv cells were isolated, expanded, activated and nucleofected with Cas9-RNPs targeting *AAVS1*, *FOXP3* and *SATB1*. The cells were cultured with or without the pro-inflammatory cytokine IL-12. **(A)** Scatter dot plot displaying KO efficiencies with median. KO efficiencies were determined by amplicon NGS sequencing and TIDE analysis. n = 2-9. **(B)** Bar graphs indicate mean of relative mRNA expression (ΔCt) of *SATB1* and *FOXP3* in *AAVS*1 and *SATB1* KO Treg cells normalized to 18S rRNA levels. RNA was isolated of FACS-sorted living cells. qPCR was performed in duplicates. n = 4-5. **(C)** Flow cytometry analysis of canonical pro- and anti-inflammatory markers in *FOXP3* KO Treg cells stimulated with or without IL-12. Percentages of marker positive cells were normalized to the respective *AAVS1* KO Treg cells with or without IL-12 stimulation. n = 6-7, ratio paired t test. **(D)** Flow cytometry gating strategy of *AAVS1* KO control Treg and Tconv cells without IL-12 conditioning. **(E)** Bar graph plots quantifying flow cytometry marker expressions of *AAVS1*, *FOXP3* and *SATB1* KO Treg cells treated with or without IL-12. Data partially also shown in Fig. 1B & 1E. n = 6-17, paired t test. **(F)** Extracellular IL-10 levels determined by LEGENDplex^TM^ assay of control-treated *AAVS1* KO and *SATB1* KO Treg cells. n = 6, paired t test. **(G)** Bar graph plots quantifying flow cytometry marker expressions of *AAVS1* and *SATB1* KO Tconv cells treated with or without IL-12. Data partially also shown in Fig. 1B &1G. n = 12-18, paired t test. **(H)** Extracellular IL-10 levels determined by LEGENDplex^TM^ assay of control-treated *AAVS1* KO and *SATB1* KO Tconv cells. n = 3, paired t test. * p<0.05, ** p<0.01, *** p<0.001, **** p<0.0001.

**Figure S2.**
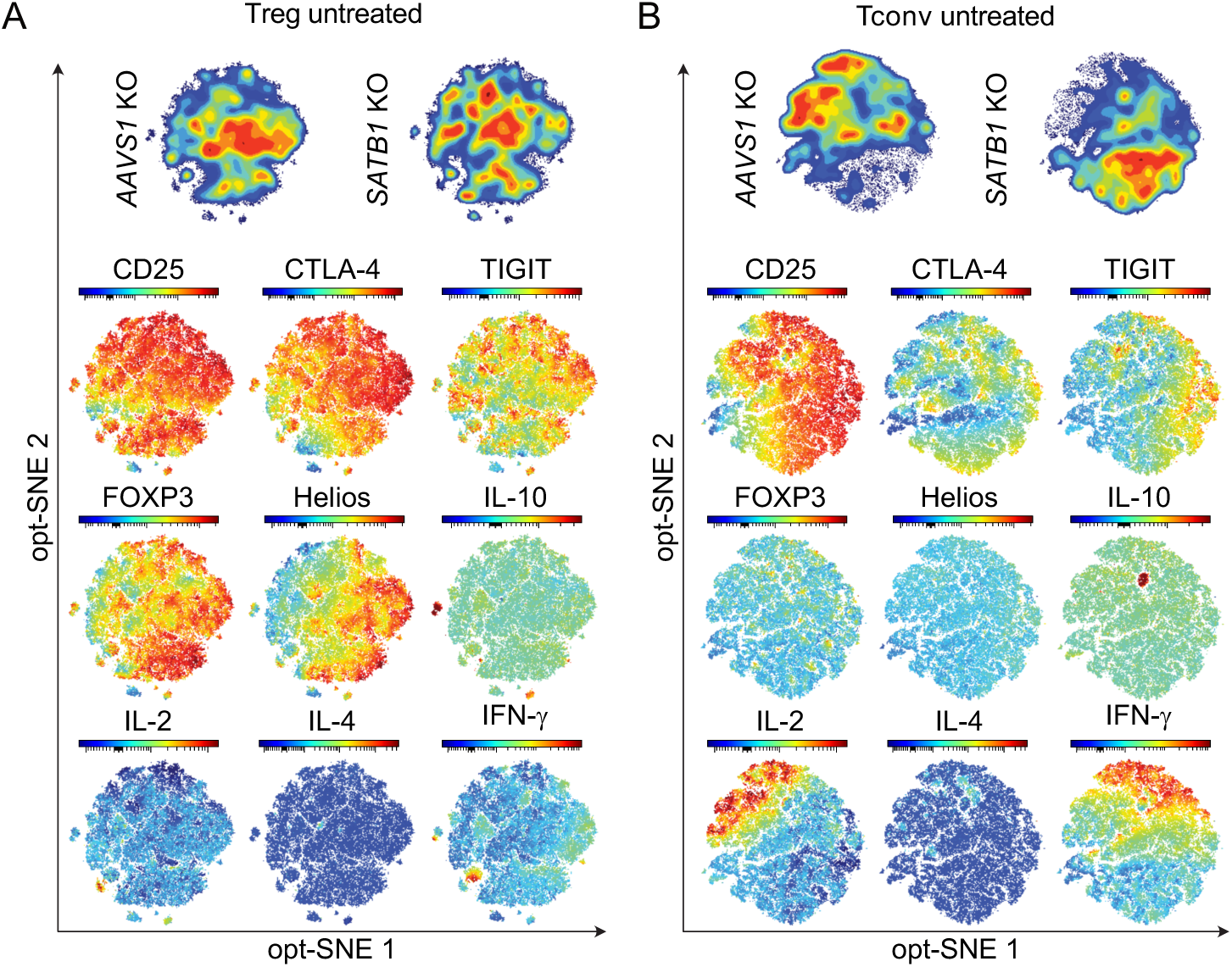
Integrated analysis of protein changes in *AAVS1* and *SATB1* KO Treg and Tconv cells based on flow cytometry. opt-SNE density plot of untreated *AAVS1* KO and *SATB1* KO Treg **(A)** and Tconv cells **(B)**. Expression levels (MFI) of tested flow cytometry markers plotted on opt-SNE plot, n(Treg) = 18, n(Tconv) = 16.

**Figure S3.**
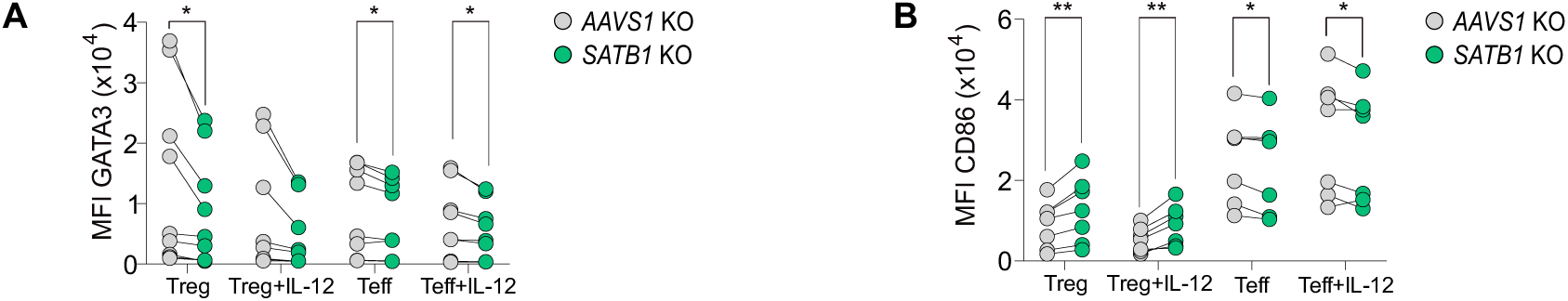
GATA3 and CD86 expression of *SATB1* KO and *AAVS1* control KO Treg and Tconv cells. **(A)** Mean fluorescence intensity (MFI) of GATA3 expression of *SATB1* KO and *AAVS1* control KO Treg and Tconv cells treated with or without IL-12. n = 6-9, paired t-test, * p<0.05. **(B)** Mean fluorescence intensity (MFI) of CD86, and FOXP3 expression of *SATB1* KO and *AAVS1* control KO Treg and Tconv cells treated with or without IL-12. n = 8-9, paired t-test, * p<0.05, ** p<0.01, *** p<0.001.

**Figure S4.**
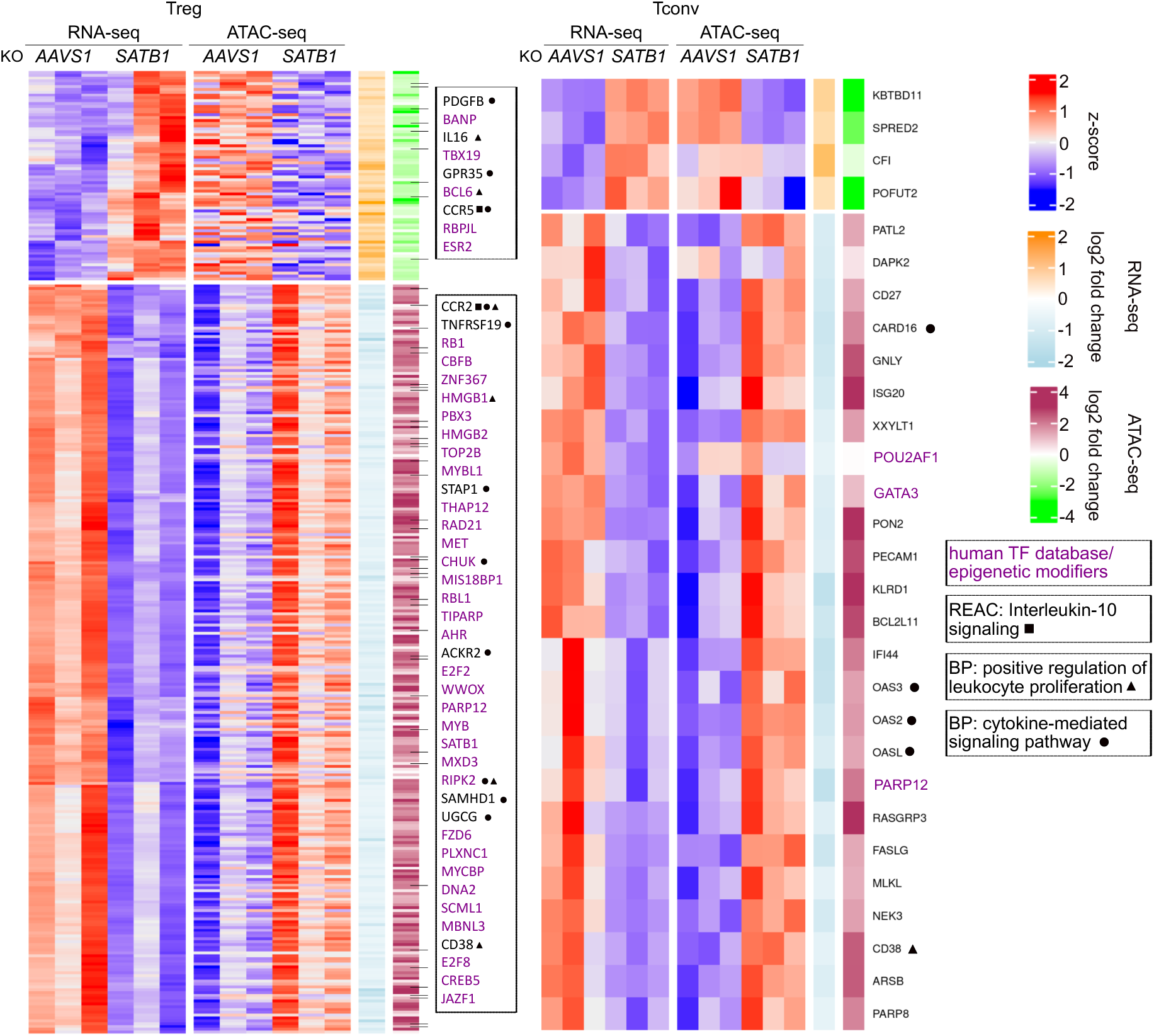
Genes differentially regulated on chromatin and transcription levels in *SATB1* KO Treg and Tconv cells. Heatmaps display z-scores of RNA- and ATAC-seq data of *SATB1* KO Treg and Tconv cells treated with IL-12. TFs differently regulated in RNA- and ATAC-seq data after *SATB1* KO are highlighted in purple. Genes associated with “Interleukin-10 signaling”, “Positive regulation of leukocyte proliferation” or “cytokine-mediated signaling” are highlighted. REAC: Reactome; BP: biological pathway.

## REFERENCES

Ahlfors H, Limaye A, Elo LL, Tuomela S, Burute M, Gottimukkala KV, Notani D, Rasool O, Galande S, Lahesmaa R (2010) SATB1 dictates expression of multiple genes including IL-5 involved in human T helper cell differentiation. Blood 116: 1443–1453

Allan SE, Crome SQ, Crellin NK, Passerini L, Steiner TS, Bacchetta R, Roncarolo MG, Levings MK (2007) Activation-induced FOXP3 in human T effector cells does not suppress proliferation or cytokine production. Int Immunol 19: 345–354

Alvarez JD, Yasui DH, Niida H, Joh T, Loh DY, Kohwi-Shigematsu T (2000) The MAR-binding protein SATB1 orchestrates temporal and spatial expression of multiple genes during T-cell development. Genes Dev 14: 521–535

Anders S, Huber W (2010) Differential expression analysis for sequence count data. Genome Biol 11: R106

Aqel SI, Yang X, Kraus EE, Song J, Farinas MF, Zhao EY, Pei W, Lovett-Racke AE, Racke MK, Li C et al (2021) A STAT3 inhibitor ameliorates CNS autoimmunity by restoring Teff:Treg balance. JCI Insight 6

Arroyo-Olarte RD, Rivera-Rugeles A, Nava-Lira E, Sanchez-Barrera A, Ledesma-Soto Y, Saavedra R, Armas-Lopez L, Terrazas LI, Avila-Moreno F, Leon-Cabrera S (2023) STAT6 controls the stability and suppressive function of regulatory T cells. Eur J Immunol 53: e2250128

Belkina AC, Ciccolella CO, Anno R, Halpert R, Spidlen J, Snyder-Cappione JE (2019) Automated optimized parameters for T-distributed stochastic neighbor embedding improve visualization and analysis of large datasets. Nat Commun 10: 5415

Beyer M, Thabet Y, Muller RU, Sadlon T, Classen S, Lahl K, Basu S, Zhou X, Bailey-Bucktrout SL, Krebs W et al (2011) Repression of the genome organizer SATB1 in regulatory T cells is required for suppressive function and inhibition of effector differentiation. Nat Immunol 12: 898–907

Bittner S, Hehlgans T, Feuerer M (2023) Engineered Treg cells as putative therapeutics against inflammatory diseases and beyond. Trends Immunol 44: 468–483

Brinkman EK, Chen T, Amendola M, van Steensel B (2014) Easy quantitative assessment of genome editing by sequence trace decomposition. Nucleic Acids Res 42: e168

Burute M, Gottimukkala K, Galande S (2012) Chromatin organizer SATB1 is an important determinant of T-cell differentiation. Immunol Cell Biol 90: 852–859

Cai S, Lee CC, Kohwi-Shigematsu T (2006) SATB1 packages densely looped, transcriptionally active chromatin for coordinated expression of cytokine genes. Nat Genet 38: 1278–1288

Chaurio RA, Anadon CM, Lee Costich T, Payne KK, Biswas S, Harro CM, Moran C, Ortiz AC, Cortina C, Rigolizzo KE et al (2022) TGF-beta-mediated silencing of genomic organizer SATB1 promotes Tfh cell differentiation and formation of intra-tumoral tertiary lymphoid structures. Immunity 55: 115–128 e119

Corces MR, Trevino AE, Hamilton EG, Greenside PG, Sinnott-Armstrong NA, Vesuna S, Satpathy AT, Rubin AJ, Montine KS, Wu B et al (2017) An improved ATAC-seq protocol reduces background and enables interrogation of frozen tissues. Nat Methods 14: 959–962

Dai A, Zhang X, Wang X, Liu G, Wang Q, Yu F (2024) Transcription factors in chimeric antigen receptor T-cell development. Hum Cell 37: 571–581

Delacher M, Simon M, Sanderink L, Hotz-Wagenblatt A, Wuttke M, Schambeck K, Schmidleithner L, Bittner S, Pant A, Ritter U et al (2021) Single-cell chromatin accessibility landscape identifies tissue repair program in human regulatory T cells. Immunity 54: 702–720 e717

Ding ZC, Shi H, Aboelella NS, Fesenkova K, Park EJ, Liu Z, Pei L, Li J, McIndoe RA, Xu H et al (2020) Persistent STAT5 activation reprograms the epigenetic landscape in CD4(+) T cells to drive polyfunctionality and antitumor immunity. Sci Immunol 5

Doan AE, Mueller KP, Chen AY, Rouin GT, Chen Y, Daniel B, Lattin J, Markovska M, Mozarsky B, Arias-Umana J et al (2024) FOXO1 is a master regulator of memory programming in CAR T cells. Nature 629: 211–218

Dominguez-Villar M, Baecher-Allan CM, Hafler DA (2011) Identification of T helper type 1-like, Foxp3+ regulatory T cells in human autoimmune disease. Nat Med 17: 673–675

Gu Z, Eils R, Schlesner M (2016) Complex heatmaps reveal patterns and correlations in multidimensional genomic data. Bioinformatics 32: 2847–2849

Gupta PK, Allocco JB, Fraipont JM, McKeague ML, Wang P, Andrade MS, McIntosh C, Chen L, Wang Y, Li Y et al (2022) Reduced Satb1 expression predisposes CD4(+) T conventional cells to Treg suppression and promotes transplant survival. Proc Natl Acad Sci U S A 119: e2205062119

Heinz S, Benner C, Spann N, Bertolino E, Lin YC, Laslo P, Cheng JX, Murre C, Singh H, Glass CK (2010) Simple combinations of lineage-determining transcription factors prime cis-regulatory elements required for macrophage and B cell identities. Mol Cell 38: 576–589

Jiang H, Lei R, Ding SW, Zhu S (2014) Skewer: a fast and accurate adapter trimmer for next-generation sequencing paired-end reads. BMC Bioinformatics 15: 182

Kakugawa K, Kojo S, Tanaka H, Seo W, Endo TA, Kitagawa Y, Muroi S, Tenno M, Yasmin N, Kohwi Y et al (2017) Essential Roles of SATB1 in Specifying T Lymphocyte Subsets. Cell Rep 19: 1176–1188

Kitagawa Y, Ohkura N, Kidani Y, Vandenbon A, Hirota K, Kawakami R, Yasuda K, Motooka D, Nakamura S, Kondo M et al (2017) Guidance of regulatory T cell development by Satb1-dependent super-enhancer establishment. Nat Immunol 18: 173–183

Köhne M, Shakiba MH, Schmidleithner L, Schulte-Schrepping J, Scholz R, Elmzzahi T, Sommer D, Li YF, Carraro C, De Domenico E et al (2025) Satb1 directs the differentiation of TH17 cells through suppression of IL-2 expression. Cell Reports 44

Lam AJ, Uday P, Gillies JK, Levings MK (2022) Helios is a marker, not a driver, of human Treg stability. Eur J Immunol 52: 75–84

Langmead B, Salzberg SL (2012) Fast gapped-read alignment with Bowtie 2. Nat Methods 9: 357–359

Laurence A, Amarnath S, Mariotti J, Kim YC, Foley J, Eckhaus M, O’Shea JJ, Fowler DH (2012) STAT3 transcription factor promotes instability of nTreg cells and limits generation of iTreg cells during acute murine graft-versus-host disease. Immunity 37: 209–222

Liao Y, Smyth GK, Shi W (2014) featureCounts: an efficient general purpose program for assigning sequence reads to genomic features. Bioinformatics 30: 923–930

Lingeman E, Jeans C, Corn JE (2017) Production of Purified CasRNPs for Efficacious Genome Editing. Curr Protoc Mol Biol 120: 31 10 31-31 10 19

Love MI, Huber W, Anders S (2014) Moderated estimation of fold change and dispersion for RNA-seq data with DESeq2. Genome Biol 15: 550

Ma H, Lu C, Ziegler J, Liu A, Sepulveda A, Okada H, Lentzsch S, Mapara MY (2011) Absence of Stat1 in donor CD4(+) T cells promotes the expansion of Tregs and reduces graft-versus-host disease in mice. J Clin Invest 121: 2554–2569

Minnar CM, Lui G, Gulley JL, Schlom J, Gameiro SR (2023) Preclinical and clinical studies of a tumor targeting IL-12 immunocytokine. Front Oncol 13: 1321318

Mirlekar B, Pylayeva-Gupta Y (2021) IL-12 Family Cytokines in Cancer and Immunotherapy. Cancers (Basel) 13

Mortazavi A, Williams BA, McCue K, Schaeffer L, Wold B (2008) Mapping and quantifying mammalian transcriptomes by RNA-Seq. Nat Methods 5: 621–628

Pallandre JR, Brillard E, Crehange G, Radlovic A, Remy-Martin JP, Saas P, Rohrlich PS, Pivot X, Ling X, Tiberghien P et al (2007) Role of STAT3 in CD4+CD25+FOXP3+ regulatory lymphocyte generation: implications in graft-versus-host disease and antitumor immunity. J Immunol 179: 7593–7604

Parkhomchuk D, Borodina T, Amstislavskiy V, Banaru M, Hallen L, Krobitsch S, Lehrach H, Soldatov A (2009) Transcriptome analysis by strand-specific sequencing of complementary DNA. Nucleic Acids Res 37: e123

Poholek AC, Jankovic D, Villarino AV, Petermann F, Hettinga A, Shouval DS, Snapper SB, Kaech SM, Brooks SR, Vahedi G et al (2016) IL-10 induces a STAT3-dependent autoregulatory loop in T(H)2 cells that promotes Blimp-1 restriction of cell expansion via antagonism of STAT5 target genes. Sci Immunol 1

Qin Z, Wang R, Hou P, Zhang Y, Yuan Q, Wang Y, Yang Y, Xu T (2024) TCR signaling induces STAT3 phosphorylation to promote TH17 cell differentiation. J Exp Med 221

Ramirez CL, Zeida A, Jara GE, Roitberg AE, Marti MA (2014) Improving Efficiency in SMD Simulations Through a Hybrid Differential Relaxation Algorithm. J Chem Theory Comput 10: 4609–4617

Schumann K, Raju SS, Lauber M, Kolb S, Shifrut E, Cortez JT, Skartsis N, Nguyen VQ, Woo JM, Roth TL (2020) Functional CRISPR dissection of gene networks controlling human regulatory T cell identity. Nature Immunology 21: 1456–1466

Stephen TL, Payne KK, Chaurio RA, Allegrezza MJ, Zhu H, Perez-Sanz J, Perales-Puchalt A, Nguyen JM, Vara-Ailor AE, Eruslanov EB et al (2017) SATB1 Expression Governs Epigenetic Repression of PD-1 in Tumor-Reactive T Cells. Immunity 46: 51–64

Sun J, Madan R, Karp CL, Braciale TJ (2009) Effector T cells control lung inflammation during acute influenza virus infection by producing IL-10. Nat Med 15: 277–284

Thakore PI, Schnell A, Huang L, Zhao M, Hou Y, Christian E, Zaghouani S, Wang C, Singh V, Singaraju A et al (2024) BACH2 regulates diversification of regulatory and proinflammatory chromatin states in T(H)17 cells. Nat Immunol 25: 1395–1410

Verstockt B, Salas A, Sands BE, Abraham C, Leibovitzh H, Neurath MF, Vande Casteele N, Alimentiv Translational Research C (2023) IL-12 and IL-23 pathway inhibition in inflammatory bowel disease. Nat Rev Gastroenterol Hepatol 20: 433–446

Yao Z, Kanno Y, Kerenyi M, Stephens G, Durant L, Watford WT, Laurence A, Robinson GW, Shevach EM, Moriggl R et al (2007) Nonredundant roles for Stat5a/b in directly regulating Foxp3. Blood 109: 4368–4375

Yasuda K, Kitagawa Y, Kawakami R, Isaka Y, Watanabe H, Kondoh G, Kohwi-Shigematsu T, Sakaguchi S, Hirota K (2019) Satb1 regulates the effector program of encephalitogenic tissue Th17 cells in chronic inflammation. Nat Commun 10: 549

Zhang Y, Liu T, Meyer CA, Eeckhoute J, Johnson DS, Bernstein BE, Nusbaum C, Myers RM, Brown M, Li W et al (2008) Model-based analysis of ChIP-Seq (MACS). Genome Biol 9: R137

